# Epigenome dysregulation resulting from NSD1 mutation in head and neck squamous cell carcinoma

**DOI:** 10.1101/2020.05.30.124057

**Authors:** Nargess Farhangdoost, Cynthia Horth, Bo Hu, Eric Bareke, Xiao Chen, Yinglu Li, Mariel Coradin, Benjamin A. Garcia, Chao Lu, Jacek Majewski

**Affiliations:** Department of Human Genetics, McGill University, Montreal, QC, H3A 1B1, Canada; McGill University Genome Centre, Montreal, QC, H3A 0G1, Canada; Department of Genetics and Development, Columbia University Irving Medical Center, New York, NY 10032, USA; Biochemistry and Molecular Biophysics Graduate Group, University of Pennsylvania, Philadelphia, PA 19104, USA; Department of Biochemistry and Biophysics, and Penn Epigenetics Institute, Perelman School of Medicine, University of Pennsylvania, Philadelphia, PA, 19104 USA

## Abstract

Epigenetic dysregulation has emerged as an important mechanism of oncogenesis. To develop targeted treatments, it is important to understand the epigenetic and transcriptomic consequences of mutations in epigenetic modifier genes. Recently, mutations in the histone methyltransferase gene NSD1 have been identified in a subset of head and neck squamous cell carcinomas (HNSCCs) – one of the most common and deadly cancers. Here, we use whole (epi)genome approaches and genome editing to dissect the downstream effects of loss of NSD1 in HNSCC. We demonstrate that NSD1 mutations are directly responsible for loss of intergenic H3K36me2 domains, followed by loss of DNA methylation, and gain of H3K27me3 in the affected genomic regions. We further show that those regions are enriched in cis-regulatory elements and that subsequent loss of H3K27ac correlates with reduced expression of their target genes. Our analysis identifies genes and pathways affected by the loss of NSD1 and paves the way to further understanding the interplay among chromatin modifications in cancer.

## INTRODUCTION

Head and neck squamous cell carcinomas (HNSCCs) are very common and deadly cancers that can develop in oropharynx, hypopharynx, larynx, nasopharynx, and oral cavity^1,2^. These anatomically^2^- and genetically^3^- heterogeneous tumors can be induced either through some behavioral risk factors—such as tobacco smoking, excessive alcohol consumption, or insufficient oral hygiene^4-7^—or through human papillomavirus (HPV)^8^ and are currently classified into HPV(-) and HPV(+) groups^9^. HPV(-) tumors constitute around 80-95% of all HNSCCs^10^. The best currently available treatments have shown promising response in HPV(+) patients but have been less successful in HPV(-) cancers^11-15^. Thus, it is of great importance to understand the etiology of HPV(-) HNSCC tumors in order to develop more effective targeted therapies.

Recently, mutations in epigenetic modifier genes, particularly the methyltransferase Nuclear Receptor Binding SET Domain Protein 1 (NSD1), have been implicated in HNSCC pathogenesis^1,16^. Subsequently, our group has identified H3K36M—*Lysine to methionine* mutations in histone H3 at the residue 36— mutations and demonstrated that NSD1 and H3K36M mutant HNSCCs form a distinct subgroup, characterized by specific DNA methylation (DNAme) patterns^17^. NSD1 is a histone lysine methyltransferase that catalyzes mono- and di-methylation of histone H3 at lysine 36 (H3K36me2)^17-20^. In addition, it may act as a transcriptional co-factor, responsible for activating or repressing different genes^21^. Mutations in other genes which encode H3K36 methyltransferases, such as NSD2 and SETD2, have not been frequently identified in HNSCC^17^, and it is not clear whether they contribute to this disease. Recent tumor-immune profiling of HNSCC patient samples has reported an association between NSD1 mutation and reduced immune infiltration^20^. In addition, it has been shown that HPV(-) tumors with NSD1 truncating mutations exhibit better treatment responses when targeted with cisplatin and carboplatin (chemotherapy based on platinum) compared to those lacking these mutations^15,22^. Thus, NSD1, its function, and the dysregulation it causes at the genetic and/or epigenetic level in HPV(-) HNSCC are of great importance for understanding the underlying mechanisms of tumorigenesis for improving the treatment responses.

We, and others, have further argued that the common mechanism underlying tumorigenicity in the H3K36me-dysregulated tumors is a reduction in H3K36me2 levels, followed by a global reduction in DNA methylation^1,17,23,24^. These observations, so far, have been based on primary tumor data, bulk quantification of epigenetic modifications, or data obtained from genetic manipulation of mouse embryonic stem cells^17,25^.

Here, we use quantitative mass spectrometry of histone post-translational modifications (PTMs), genome-wide Chromatin Immunoprecipitation Sequencing (ChIP-Seq), and Whole Genome Bisulfite Sequencing (WGBS) to finely characterize the differences in epigenetic characteristics of NSD1-Wildtype (NSD1-WT) and NSD1-Mutant (NSD1-MT) HNSCC cell lines. Next, we utilized CRISPR-Cas9 genome editing to inactivate NSD1 in several independent cell lines and showed that, in an isogenic context, the ablation of NSD1 is sufficient to recapitulate the epigenetic patterns that were observed in the patient-derived material. Furthermore, carried out RNA sequencing and characterized the transcriptional impact of NSD1 loss in order to link epigenetic programming with functional outputs. We directly demonstrate the connection between NSD1, H3K36me2, Polycomb Repressive Complex 2 (PRC2)-mediated H3K27me3, and DNA methylation modifications in HNSCC. We also link the depletion of intergenic H3K36me2 with reduced activity of cis-regulatory elements– as profiled by the levels of H3K27ac – and reduced levels of expression of target genes.

## RESULTS

### Epigenomic Characterization of NSD1 WT and Mutant HNSCC cell lines

We have previously shown that H3K36M and NSD1 mutations in HNSCCs are associated with low global levels of H3K36me2^17^. More recently, we provided evidence that NSD1 mutant HNSCC samples are specifically characterized by low *intergenic* levels of H3K36me2^25^. To confirm those results in a larger number of samples and characterize additional epigenetic marks, we examined three NSD1-WT (Cal27, FaDu, Detroit562) and three NSD1-MT (SKN3, SCC4, BICR78) patient-derived HNSCC cell lines (Supplementary Table 1). Mass spectrometry analysis demonstrates a clear difference in the global levels of H3K36me2 when comparing the mean of NSD1-WT with the mean of NSD1-MT samples (Fig. 1a, Supplementary Data 1). Visualization of H3K36me2 ChIP-seq tracks in representative regions illustrates that, in NSD1-MT cell lines, this mark is significantly reduced at the intergenic regions adjacent to genic regions (Fig. 1b). This intergenic depletion of H3K36me2 can be generalized to a genome-wide scale using heat maps and box plots (Fig. 1c, d). We note that there is significant variability across NSD1-WT cell lines with respect to the distribution of intergenic H3K36me2 (Fig. 1c): in FaDu, nearly all intergenic regions are marked with H3K36me2, and as a result, intergenic levels are higher compared to genic, while Cal27 has the least pronounced intergenic H3K36me2 domains. This is further clarified with each cell line being illustrated individually (Supplementary Fig. 1). Those differences are likely to be an effect of the cell of origin, presence of other oncogenic mutations, and relative activity levels of epigenetic modifier enzymes. However, our analysis shows a consistent and nearly total lack of intergenic H3K36me2 in all NSD1-MT cell lines, in contrast to genic levels that remain comparable to NSD1-WT lines.

**Fig. 1.**
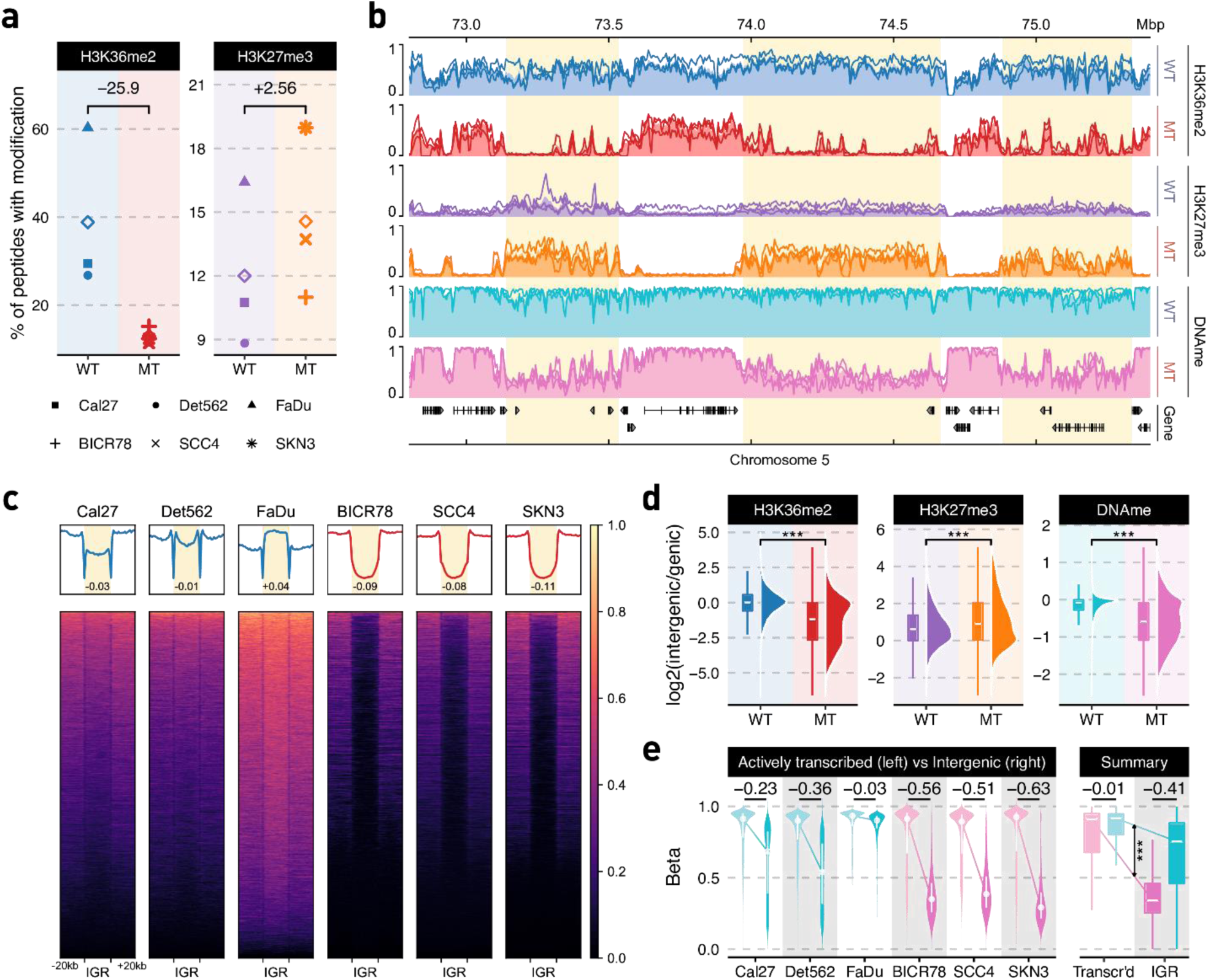
Epigenomic characterization of NSD1-WT and mutant HNSCC cell lines. **a** Genome-wide prevalence of modifications based on mass spectrometry; diamonds represent within-condition averages; p-values obtained using Welch’s t-test: H3K36me2 p=0.07 and H3K27me3 p=0.24. **b** Genome-browser tracks displaying individual samples as lines and condition averages as area plots in a lighter shade; ChIP-seq signals shown are MS-normalized logCPM while beta values are used for WGBS; regions of noticeable difference are highlighted. **c** Heatmaps showing H3K36me2 (MS-normalized logCPM) enrichment patterns +/− 20kb flanking intergenic regions (IGR). Number displayed at the bottom of aggregate plots correspond to the intergenic / genic ratio where TSS/TES and outer edges are excluded. **d** Relative enrichment of signal within intergenic regions over those of flanking genes; CPM values are used for ChIP-seq and beta values for WGBS; *** = Wilcoxon rank sum test p-value < 1e-5; **e** Distribution of DNA methylation beta values within actively transcribed genes (zFPKM^27^ > −3) compared against those in intergenic regions.

We have previously observed that NSD1 and H3K36M mutant HNSCC tumors are hypomethylated at the DNA level^17^ and, using mouse cell line models, proposed that this hypomethylation is mechanistically linked to the decrease in intergenic H3K36me2 via reduced recruitment of the *de novo* DNA methyltransferase, DNMT3A^25^. Our extended analysis here clearly indicates that the decrease of intergenic H3K36me2, corresponds to a significant decrease in intergenic DNA methylation in all three NSD1-MT, as compared to NSD1-WT HNSCC cell lines (Fig. 1b, e). Concurrently, DNA methylation levels within actively transcribed genes remain comparable across all profiled cell lines, irrespective of their NSD1 status. Again, we note considerable variability across cell lines, with FaDu, which has the highest levels of global and intergenic H3K36me2, possessing a globally hypermethylated genome.

Finally, we examined the silencing mark H3K27me3^26^, since its levels and distribution have been shown to be negatively correlated with H3K36me2. Mass spectrometry shows an elevated level of H3K27me3 in the NSD1-MT cell lines (Fig. 1a). Through ChIP-seq of H3K27me3 we observe that it is the intergenic regions depleted of H3K36me2 in all 3 NSD1-MT samples that specifically exhibit a corresponding increase in H3K27me3, corroborating the antagonistic relation between these two marks (Fig. 1d, Supplementary Fig. 2). Overall, our observations demonstrate that lack of intergenic H3K36me2 that characterizes NSD1-MT HNSCC samples is associated with decreased intergenic DNA methylation levels and increased H3K27me3 levels in HPV(-) HNSCC.

### Knock-out of NSD1 is sufficient to recapitulate the decrease in intergenic H3K36me2 and confirms the relationship with DNA methylation and H3K27me3

In order to demonstrate that our observations in patient-derived material are a direct consequence of the presence or absence of NSD1 mutations, we used the CRISPR-Cas9 system to edit the three NSD1-WT HNSCC cell lines and generate several independent NSD1-knockout (NSD1-KO) clonal cultures per cell line. This approach ensures an isogenic context; that is, we can isolate the effect of NSD1 by deleting it on an otherwise unaltered genetic background. Using three different cell lines generalizes the results across genetic backgrounds. Propagation of multiple independent clones for each parental line minimizes the possible off-target and clone-specific effects. We targeted the SET domain and PWWP domain of NSD1 since these two domains play crucial roles in catalyzing the deposition of methyl groups to H3K36^28^ and reading the methylated lysines on Histone H3^29^, respectively. Thus, disruption of either or both of these domains by CRISPR-Cas9 is likely to compromise the function of NSD1 as a histone methyltransferase. We successfully generated three HNSCC isogenic NSD1-KO clones in Detroit562, two in Cal27, and one in the FaDu cell line. The editing was confirmed by sequencing (MiSeq) of the regions surrounding the target sites (Supplementary Fig. 3, Supplementary Table 2) and the absence of the protein was confirmed by Western blots of NSD1 (Supplementary Fig. 4). Mass spectrometry and ChIP-seq were used to quantify genome-wide levels and distribution of H3K36me2 and H3K27me3 in NSD1-KO HNSCC isogenic cell lines and all their replicate clones compared to their parental cell lines. Similar to our observations comparing primary NSD1-MT and NSD1-WT HPV(-) HNSCC cell lines, NSD1-KO isogenic cell lines show a global reduction of H3K36me2 and an increase in H3K27me3 levels (Fig. 2a, Supplementary Data 1) most prominently in intergenic regions (Fig 2b, Supplementary Fig. 5). Using mass spectrometry-normalized ChIP-seq data, we emphasized the pronounced pattern of H3K36me2 depletion and H3K27me3 enrichment at intergenic regions (Fig. 2c, Supplementary Fig. 6), the degree of which can be further quantified by comparing intergenic enrichment levels against flanking genes (Supplementary Fig. 7). We, next, profiled the genome-wide DNA methylation changes resulting from NSD1-KO. DNA methylation was decreased predominantly in intergenic regions, and this reduction was particularly pronounced in regions that lost H3K36me2 (Fig. 2d). Firstly, we note that the degree of DNAme reduction varied across cell lines: FaDu, which in the parental cell lines had a globally hypermethylated genome, exhibited the highest (8.1%), Cal27 showed an intermediate (3.7%), and Detroit562 showed the lowest (2.3%) DNAme reduction in regions losing H3K36me2 (Fig. 2d). While the results are consistent with the previous observation that H3K36me2 recruits active DNA methyltransferases^25^, it also demonstrates that other factors, which are likely to be dependent on the genetic and epigenetic states of the parental cells, also play important roles. In the case of DNA methylation, those may include the relative importance of maintenance, versus *de novo*, DNA methyltransferases^30^, or levels of relevant metabolites, such as SAM^31^. Secondly, we observe that within active genes DNA methylation levels remain nearly unchanged in NSD1-KO, and actually exhibit a slight increase (Fig. 2d). This suggests that the presence of H3K36 methylation within actively transcribed genes is still sufficient to maintain DNAme in those regions. Finally, we find that the reduction in DNAme, while consistent across NSD1-KO cell lines, individual clones, and statistically significant, is considerably smaller than the difference observed between NSD1-WT and NSD1-MT patient derived cell lines. Overall, the extent to which NSD1-KOs recapitulate the epigenetic characteristics of NSD1-MT cell lines is highest for H3K36me2, intermediate for H3K27me3, and lowest for DNAme (Fig. 2e). To validate the reproducibility across several independent knockouts generated from different cell lines, we specifically examined the differential levels of H3K36me2, H3K27me3, and DNA methylation for parental (wild-type) versus knockout (PA-KO) and wild-type versus mutant (WT – MT) comparisons. We expectedly observe in both cases a strong correlation between H3K36me2 and DNAme, which contrasted by the antagonism between H3K36me2 and H3K27me3 (Supplementary Fig. 8). Moreover, we find that despite notable variation among different cell lines, knockouts of the same cell lines are highly correlated – indicating a high degree of consistency in terms of the effect exerted by NSD1-KO among all three epigenetic modifications (Supplementary Fig. 9).

**Fig. 2.**
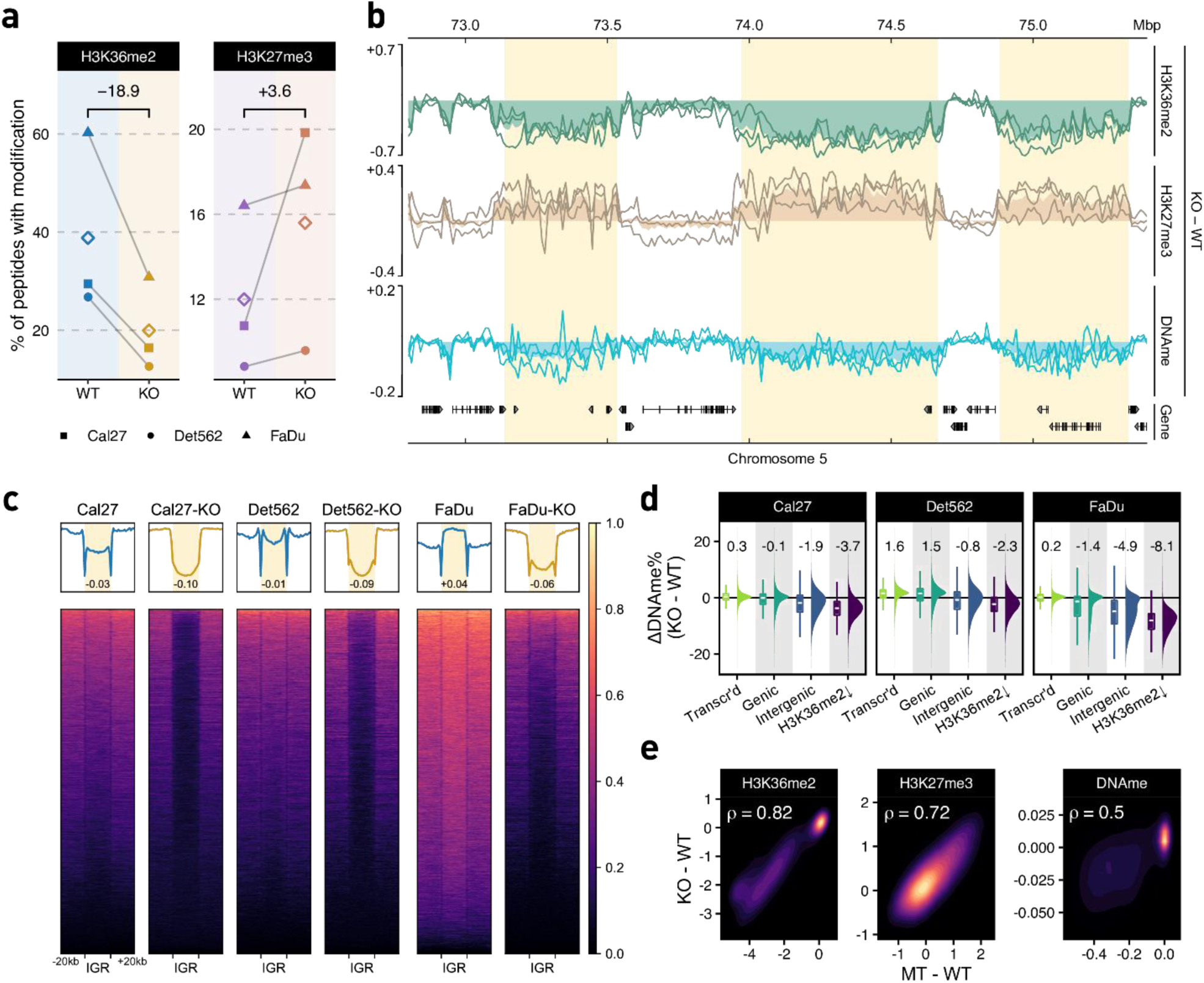
Epigenomic characterization of NSD1-WT and KO HNSCC cell lines. **a** Genome-wide prevalence of modifications based on mass spectrometry; diamonds represent within-condition averages; p-values obtained using Welch’s t-test: H3K36me2 p=0.04, H3K27me3 p=0.16. **b** Genome-browser tracks displaying individual cell-line differences (KO-PA) as lines and condition averages as area plots in a lighter shade; ChIP-seq signals shown are MS-normalized logCPM while beta values are used for WGBS; regions of noticeable difference in Fig. 1b are highlighted. **c** Heatmaps showing H3K36me2 enrichment patterns centered on intergenic regions (IGR). Number displayed at the bottom of aggregate plots correspond to the intergenic / genic ratio where TSS/TES and outer edges are excluded. **d** Distribution of differential beta values within actively transcribed genes, all genes, intergenic regions, and regions depleted of H3K36me2 (corresponding to regions defined in Fig. 3a as “cluster B”); median values are shown at the top. **e** Spearman correlation of differential enrichment between NSD1-WT vs KO and WT vs MT.

### Loss of NSD1 preferentially impacts intergenic regulatory elements

It is of paramount interest to understand the downstream functional consequences of the epigenetic remodeling from NSD1’s deletion. To identify and further characterize the genomic compartments that exhibit the highest loss of H3K36me2, we subdivided the genome into 10kb bins and compared each parental NSD1-WT line with its respective NSD1-KO clones (Fig. 3a, Supplementary Fig. 10), showing that H3K36me2 profiles of genomic bins subdivide into three distinct clusters. The lower left cluster (A) corresponds to regions with negligible levels of H3K36me2 in both WT and KO. The upper right cluster (C) contains regions that maintain high H3K36me2 levels under both conditions; those regions are predominantly genic (color-coded in blue). The lower right quadrant (cluster B) represents mostly intergenic regions (color-coded red) with high initial levels of H3K36me2 in the parental lines and low levels in the knockout. Examination of gene expression data revealed that the few genes overlapping cluster B bins were lowly expressed across all samples, and thus resemble intergenic regions at the transcriptional level (Supplementary Fig. 11). We used genomic element annotations (Ensembl Regulatory Build^32^) and carried out enrichment analysis to compare the regions affected to those not affected by the loss of H3K36me2. Among the strongest observed functional enrichment categories in H3K36me2 regions (cluster B) were cis-regulatory elements^33^ (CREs): promoter flanking regions and enhancers (Fig. 3b, Supplementary Data 2). We also observed an enrichment in CTCF binding sites, suggesting that the regions that lose H3K36me2 are enriched in chromosomal contacts that are characteristic of transcriptionally active regions^34^. In contrast, we note that bins in cluster A, the low-invariant H3K36me2 regions, were only associated with the broadly defined intergenic classification and did not exhibit an enrichment of any annotated regulatory categories (Supplementary Fig. 12, Supplementary Data 2). The reduction of H3K36me2 at the CREs was accompanied by a corresponding decrease in DNA methylation and an increase in the silencing mark H3K27me3 (Fig. 3c). However, the increase in H3K27me3 appeared to be much more focused, as seen from the narrower peak width, suggesting that the gain of H3K27me3 is specific to these elements rather than across the entire region experiencing a loss of H3K36me2. Finally, the CREs located in the regions depleted of H3K36me2 experience a sharp reduction, which is highly specific to the location of the CRE, of the active chromatin mark H3K27ac (Fig. 3c). These results suggest that intergenic regions that are affected by the deletion of NSD1 and subsequent loss of H3K36me2 exhibit a reduced regulatory potential. Although mass spectrometry shows that global levels of H3K27ac appear to increase between NSD1-WT and NSD1-KO (Supplementary Fig. 13, Supplementary Data 1), focusing on bins that specifically lose H3K36me2 in NSD1-KO cells (Cluster B), we observed almost exclusively peaks with reduced H3K27ac binding (Fig. 3d). Irrespective of H3K36me2 changes, peaks that gain H3K27ac in NSD1-KO showed a genomic distribution resembling the set of all consensus peaks, whereas those with reduced intensities were preferentially located in distal intergenic regions (Fig. 3e). Using transcription start sites (TSS) as a reference point, we reach a similar conclusion, finding that down-regulated H3K27ac peaks are preferentially located away from the TSS (Supplementary Fig. 14).

**Fig. 3.**
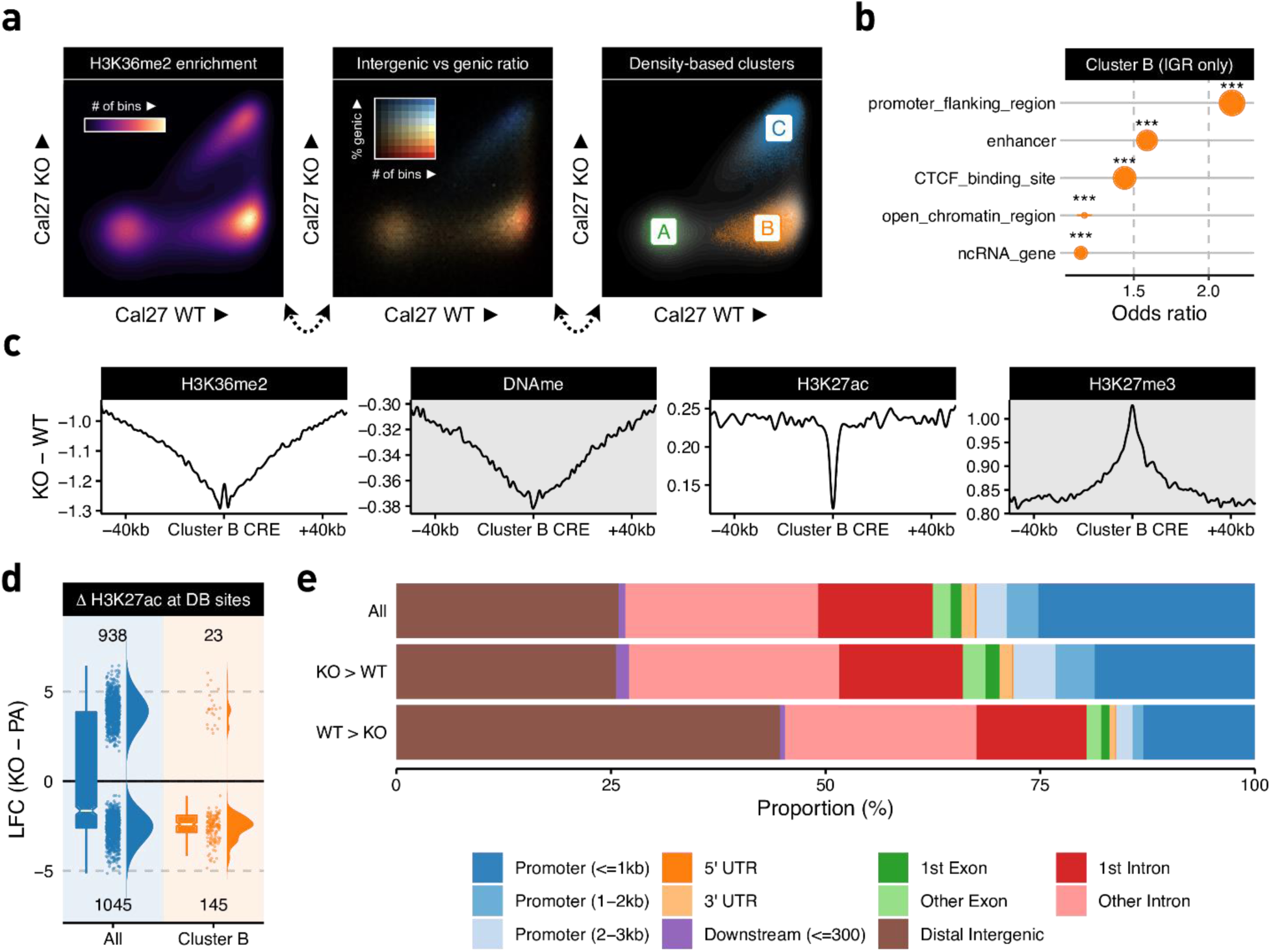
Loss of NSD1 preferentially impacts distal intergenic cis-regulatory elements **a** Scatterplots of H3K36me2 enrichment (10kb resolution) comparing a representative WT parental sample (Cal27) against its NSD1-KO counterpart (see Supplementary Fig. 7 for other cell lines) **b** Overlap enrichment result of Ensembl annotations with bins consistently labelled as cluster B (i.e., identified as B in all three paired WT vs KO comparisons). Stratification is applied to only focus on intergenic regions to avoid spurious associations to annotations confounded by their predominantly intergenic localization. The size of the dots corresponds to number of bins overlapping the corresponding annotation. **c** Aggregate plots of differential signal enrichment centered around CREs overlapping consistent cluster B bins. Values are averaged across all three WT vs KO comparisons. **d** Log fold change of H3K27ac normalized enrichment values comparing all differentially bound sites to those overlapping consistent cluster B bins **e** Distribution of genomic compartments overlapping various subsets of H3K27ac peaks categorized by differential binding status.

### Loss of H3K36me2 domains and enhancer H3K27ac affects the expression of target genes

We next investigated how the loss of NSD1-mediated intergenic H3K36me2 affects transcriptional activity, by comparing gene expression between the NSD1-WT and NSD1-KO cells. Overall, we did not find a large imbalance between upregulated and downregulated genes, with slightly more genes significantly increasing (179) than decreasing (145) expression in NSD1-KO lines (Fig. 4a, Supplementary Fig. 15a). Epigenetic dysregulation generally results in massive transcriptional changes^35-37^, many of which may represent downstream effects and not be directly related to the effect of the primary insult. Based on our analysis of H3K27 acetylation, we hypothesized that the primary reduction in gene activity is mediated by the change of the epigenetic state of CREs. Hence, we next investigated the specific effect of H3K36me2 loss at enhancers on the expression of their predicted target genes. We used a high-confidence set (“double-elite”) of pairings obtained from GeneHancer^38^ to associate distal epigenetic changes to putative target genes. By comparison to all DEGs, those targeted by CREs depleted of H3K36me2 are mostly downregulated in NSD1-KO (43 down versus 6 up, Fig. 4a). We expanded the analysis by directly considering the H3K27ac states of all annotated enhancers and subdividing them into three subsets: significantly increased, not significantly changed, and significantly decreased in NSD1-KO. We found that genes paired with enhancers exhibiting reduced acetylation undergo a relatively large decrease in expression (median LFC = −0.695), as compared to the upregulation of genes whose enhancers increase in acetylation (median LFC = 0.233, Fig. 4b). We conclude that the reduction in enhancer activity has a dominant effect on the regulatory landscape of NSD1-KO cells, in comparison to the non-H3K36me2 associated increase in acetylation.

**Fig. 4.**
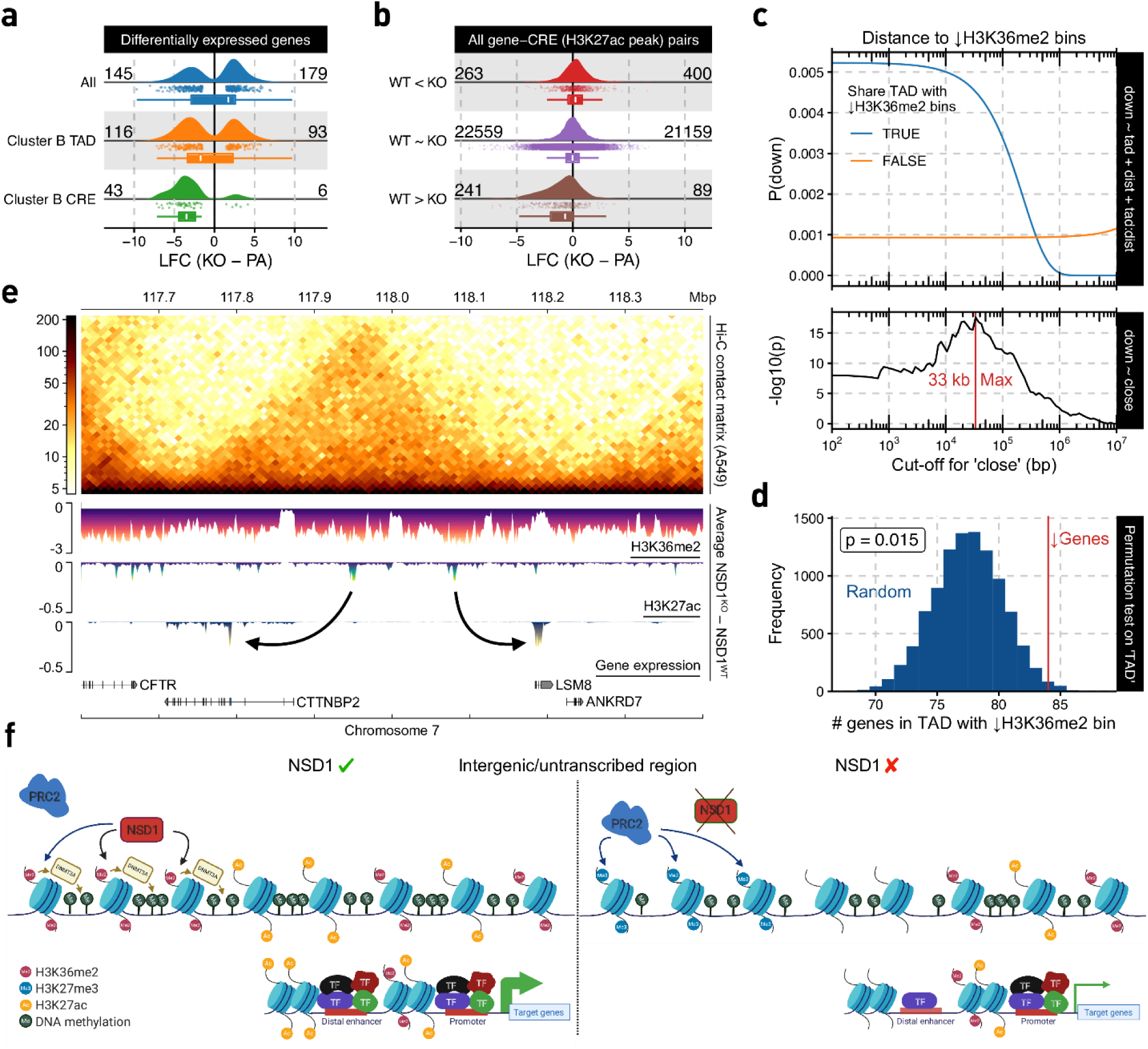
Loss of H3K36me2 domains and enhancer H3K27ac affects expression of target genes. **a** Log fold-change (LFC) of various subsets of DEGs. **b** LFC of putative target genes for various differential binding (DB) site subsets. **c** Logistic regression model outputs for expression down-regulation status based on its distance to and/or whether it shares a TAD with a cluster B bin. **d** Permutation test on down-regulated genes’ tendency to share a TAD with cluster B bin, controlling for distance. **e** Example loci illustrating genome-wide phenomenon using differential signal tracks in which enrichment values of the respective parental line were subtracted from the corresponding knockouts, after which the average across lines was taken. **f** Schematic model of epigenetic dysregulation resulting from the absence of NSD1 (created with BioRender.com). Note that in the absence of NSD1, PRC2 deposits H3K27me3 in the same intergenic regions where H3K36me2 was depleted. In addition, H3K27ac decreases around distal enhancers located in these H3K36me2-depleted regions.

Since it was recently proposed that the effect of intergenic epigenetic changes is constrained by chromatin conformation^39,40^, we considered the effect of H3K36me2 depletion at the level of Topologically Associated Domains (TAD). We extracted a publicly available TAD information from an epithelial lung cancer cell line A549^41^, which we expect to have comparable chromatin conformation to epithelial HNSCC^1,41-44^. Specifically, we investigated if a decrease in gene expression is over-represented in TADs containing H3K36me2-depleted regions (Fig. 3a, cluster B). As an association could also arise due to simple linear — rather than spatial — proximity, we included the distance of genes to their nearest cluster B bin as a covariate together with the status of sharing a TAD in logistic regression modeling (Fig. 4c, Supplementary Data 3). While the likelihood of reduced expression remains low when TAD boundaries fence-off genes from their closest H3K36me2-depleted regions, in the absence of such elements a strong association (p-value of ‘TAD’ = 5e-8) that decays with distance can be observed. Next, in order to exclude the possibility that we may not be appropriately accounting for the distance effect, the continuous distance variable was substituted with a categorical surrogate to signify whether or not a gene is within a critical distance of the nearest cluster B bin. Subsequent to selecting the distance cut-off (at 33 kb) that exhibited the strongest association with lowered expression, we found that the presence of H3K36me2-depletion in the same TAD remained a significant contributing factor (p = 0.02). Finally, in an alternative approach to control for the distance effect, we used resampling to create lists of genes that are equal in size to the set of down-regulated genes and have the same distribution of distance to the nearest cluster B bin. Again, we found that in this controlled comparison there was still a significant tendency for down-regulated genes to occupy the same TAD as H3K36me2 depleted regions (Fig. 4d).

We conclude that, as has been suggested in other systems^39,40^, that the effects of H3K36me2 depletion in HNSCC cells are governed in part by 3D chromatin structure. We also propose that the distance of 36 kb may represent an average distance between a depleted enhancer and its target gene.

Overall, we show that the loss of intergenic H3K36me2 domains in NSD1-KO cell lines results in loss of H3K27ac and enhancer activity of the affected regions, leading to a reduction in expression of target genes, and that this effect is more significant within a surrounding TADs than outside of the TAD (see Fig. 4e for a representative chromosomal region). To summarize our data, we generated a schematic model of epigenome dysregulation resulting from the absence of NSD1 (Fig. 4f). Upon the knockout of NSD1, intergenic H3K36me2 levels drop significantly, H3K27me3 increases in the same regions and DNAme slightly decreases around those regions that are depleted of H3K36me2. In addition, at those H3K36me2-depleted regions, H3K27ac decreases, primarily at distal enhancers, making those enhancers weaker/less active. These changes in the strength of distal enhancers will consequently lead to lower expression of the genes that they regulate.

### Transcriptomic changes and pathways affected by the absence or loss of NSD1 and H3K36me2

Having established that the loss of NSD1-mediated H3K36me2 specifically affects cis-regulatory elements, resulting in concomitant decreases of H3K27ac and expression of putative target genes (Fig. 4c, Supplementary Data 3), we set out to characterize the downstream transcriptomic alterations. We first focused on the primary targets of NSD1 deletion, i.e. the genes directly affected by the loss of intergenic H3K36me2. Those are most likely to resemble the early oncogenic events that occur in the cell of origin of NSD1-MT tumors. We took an integrative approach by jointly evaluating H3K36me2, H3K27ac, and RNA-seq data. GeneHancer^38^ links residing within the same TAD (projected from the A549 cell line data^41^) as regions depleted of H3K36me2 (Fig. 3a, cluster B bins) were filtered for CREs overlapping our merged H3K27ac peak-set while also presenting changes in agreement with those of gene expression. The final set of ∼5000 pairs was ordered independently by the differential test statistics for each assay type, after which a single ranking was obtained through taking the geometric mean, enabling us to perform gene set enrichment analysis (GSEA)^45,46^. This approach is valuable in synthesizing information and extracting biological meaning from long lists of differentially expressed genes. Several representative “gene sets” have been established to date. Here, we highlight the analysis using the “Hallmark” gene sets^47^, which we found to efficiently condense redundant annotations while still retaining the main trends observed in the data. Five of the seven significantly overrepresented gene sets were associated with decreased activity in the NSD1-KO condition (Fig. 5a, Supplementary Table 3). Most of these molecular signatures are consistent with previously reported roles of NSD1/H3K36me2 in immune response^20^, epithelial-mesenchymal transition (EMT)^48-51^, and regulation of RAS signaling^52,53^. Our analysis suggests that NSD1 mutations may facilitate HNSCC development through its pleiotropic effects on tumor immunity, signaling, and plasticity.

**Fig. 5.**
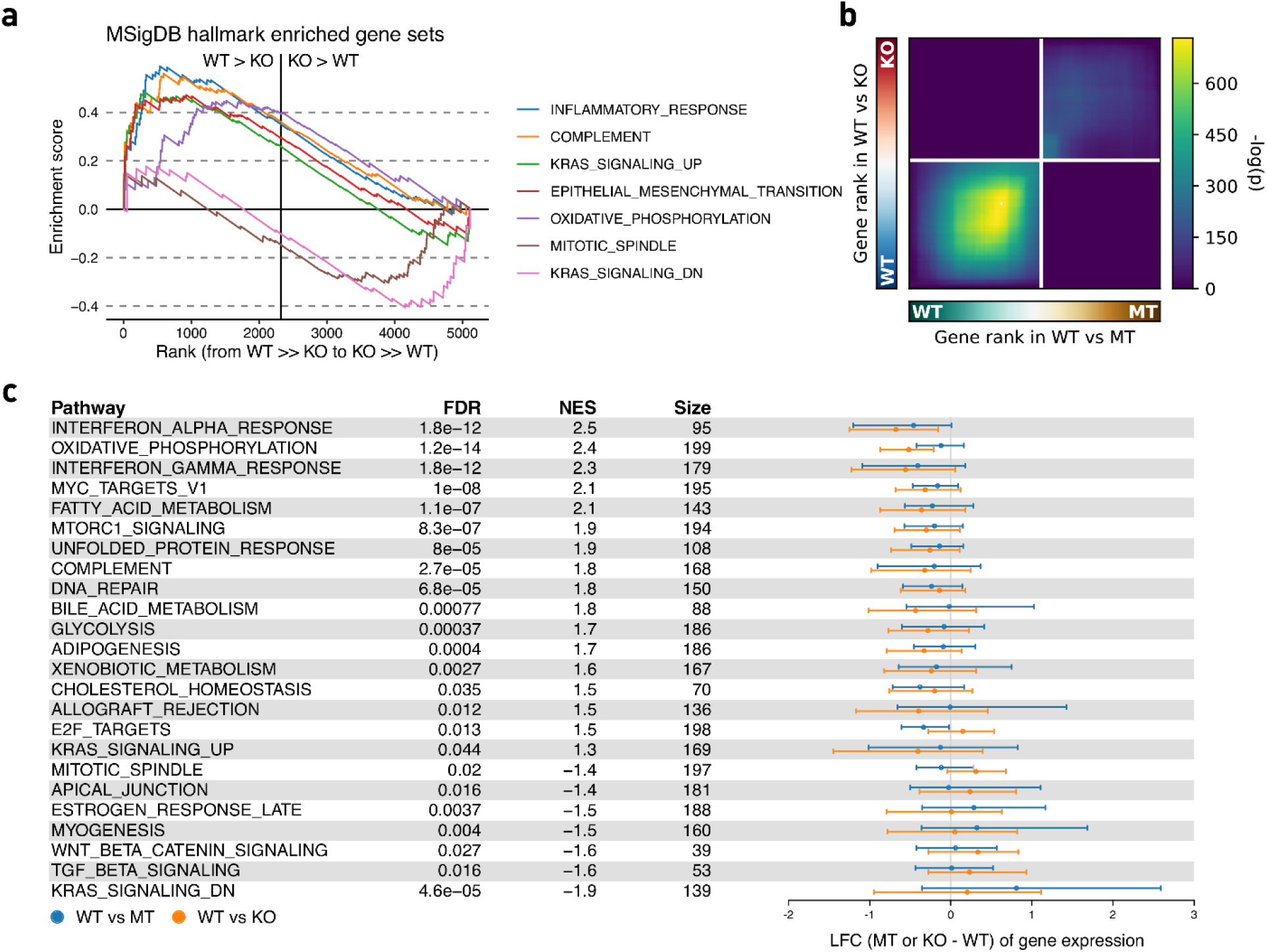
Changes in transcriptome and pathways resulted from loss of NSD1 and reduced H3K36me2 levels. **a** GSEA enrichment plot of hallmark gene sets significantly associated with the aggregated ranking of differentially expressed genes and genes targeted by differentially acetylated enhancers using their test statistics. **b** Stratified rank-rank hypergeometric overlap plot of gene expression differences between NSD1-WT vs MT and WT vs KO. **c** Distribution of expression changes for leading edge genes of hallmark gene sets significantly associated with the aggregated ranking of differential gene expression for both NSD1-WT vs MT and WT vs KO.

Finally, we wanted to connect the transcriptional characteristics of NSD1-KO HNSCC cell lines to those observed in patient-derived NSD1-MT cell lines. We reasoned that identifying the overlap between those two sets may help correct for the cell of origin and other confounding factors and highlight the pathways downstream of NSD1 deletion. Although we observed a propensity for up-regulation in both contrasts (Supplementary Fig. 15a, b), we found a significant overlap for down-regulated genes – to a degree much stronger than between those on the contrary – using a fixed p-value cut-off. By relaxing the criteria and using a rank-rank hypergeometric overlap^54^, we illustrate the extent of the concordance and also show that it is most pronounced in genes that experience downregulation (but not upregulation) in the absence of NSD1, while we find no enrichment in discordant expressional changes (Fig. 5b). GSEA using the Hallmark gene sets again identifies interferon (alpha and gamma) response to be among the top pathways that are downregulated in the absence of NSD1. Several other processes, such as oxidative phosphorylation and metabolism are also notably affected (Fig. 5c, Supplementary Table 4).

### Validation of cell line-based observations in primary tumors from TCGA

We additionally carried out a detailed re-analysis of primary tumor data from TCGA-HNSC to validate the results obtained from our simple knockout system. By first ranking patient samples based on their relative similarity to our cell lines (i.e., greater resemblance to NSD1-WT or to KO), in terms of either their methylome and transcriptome, we found that tumors most similar to NSD1-KO cell lines are highly enriched for damaging NSD1 alterations (Fig. 6a). Upon closer examination of the DNA methylation landscape in these tumors, we not only recover previous findings that DNA hypomethylation in NSD1-mutants preferentially affects CREs (Supplementary Fig. 16) but also find that the degree of DNA hypomethylation is significantly associated to their overlap with regions depleted of H3K36me2 upon NSD1-KO previously discussed (Fig. 6b). Furthermore, we observed that as with the NSD1-WT vs MT and vs KO cell line expressional comparison (Fig. 5d), contrasting differential gene expression of NSD1-WT vs KO in cell lines against NSD1-WT vs MT tumors also revealed a similarly striking concordance of down-regulation (Fig. 6c).

**Fig. 6.**
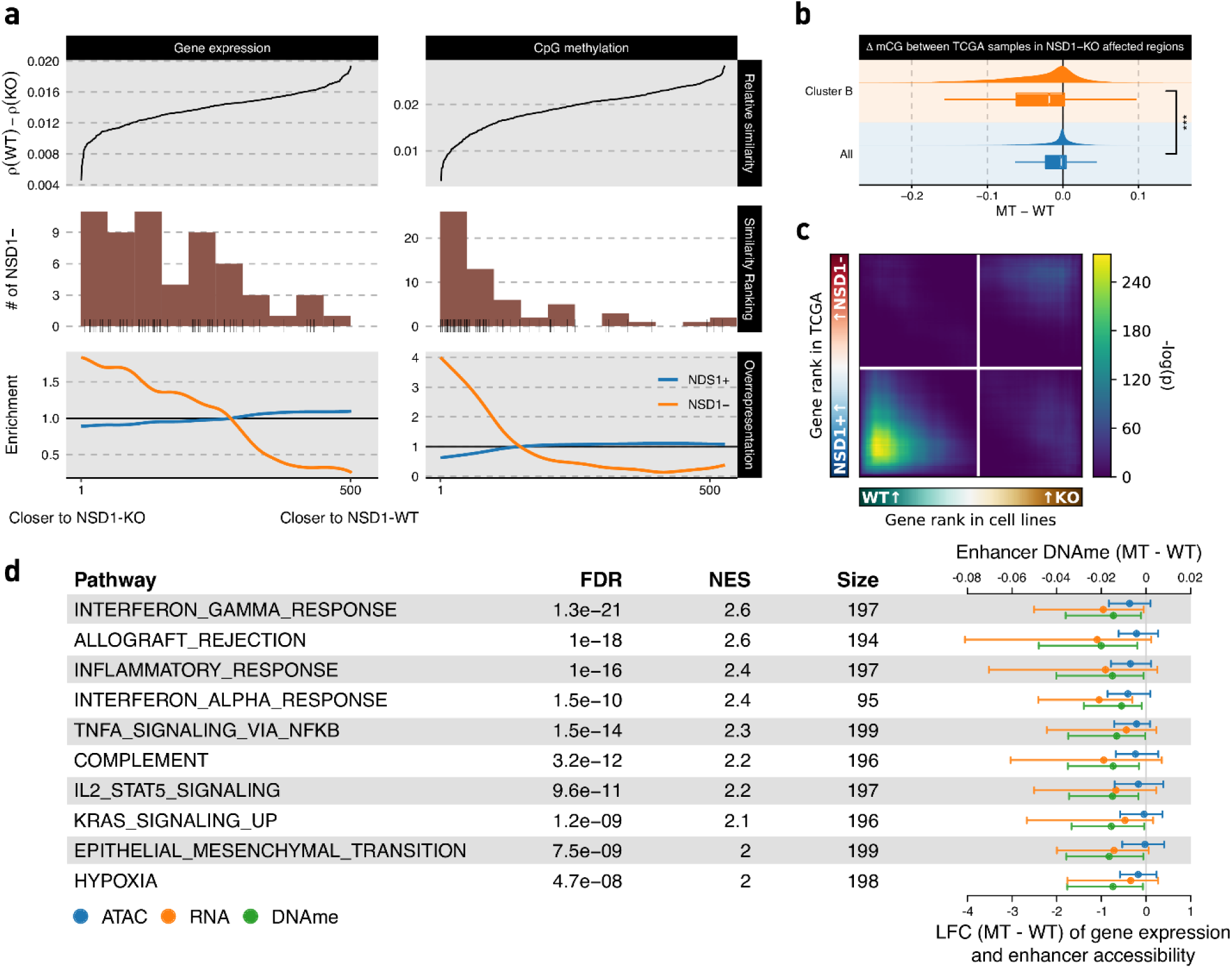
Validation of cell line-based observations in primary tumors from TCGA. **a** TCGA-HNSC samples were ranked by relative similarity to NSD1-WT and -KO cell line samples (top), NSD1 mutational status were tabulated (middle), and enrichment of NSD1 mutational groups within quantiles was computed (bottom); **b** Differential CpG methylation across all sites are contrasted against those located in regions depleted of H3K36me2 upon NSD1-KO (i.e., cluster B bins); **c** Rank-rank hypergeometric overlap of genes ranked by LFC in cell line system and TCGA, see 5d. **d** Most significant results from GSEA on genes ranked by concordant up-regulation (or down-regulation) of gene expression and enhancer accessibility and DNA methylation, with the values associated with constituent genes shown to the right.

Having established that key differences between NSD1-WT and KO cell lines largely capture the dichotomy between primary NSD1+/− tumors in various genomic modalities independently, we proceeded to adopt an integrative framework. Specifically, we performed rank aggregation across three lists of genes ranked by differential expression, differential enhancer methylation, and differential enhancer accessibility in order to identify genes which are repressed upon decreased distal regulatory potential and vice versa. As a result, we find that among hallmark gene pathways “interferon response” and other immune pathways as well as “epithelial-mesenchymal transition” are once again exceptionally significant with positive enrichment scores, signifying that genes within these pathways are expressed at lower levels and have less active cis-regulatory elements in the absence of functional NSD1 (Fig. 6d).

## DISCUSSION

HPV(-) HNSCC is a deadly cancer^55,56^ and, despite the use of innovative targeted and immune-therapies, treatments have not been effective, primarily due to poor understanding of the underlying tumorigenesis mechanisms^11,15,56^. We have previously identified a subset of HPV(-) HNSCCs that is characterized by loss of function NSD1 or H3K36M mutations with unique molecular features^17^. More recently, NSD1 has been demonstrated to be a potential prognostic biomarker in HPV(-) HNSCC^15^, suggesting that a distinct biological mechanism is involved during the evolution of NSD1-mutant HNSCC. Thus, in order to improve treatment outcomes, we need to understand how NSD1 mutations contribute to the formation or progression of this cancer.

Our comparison of three patient-derived NSD1-WT and three NSD1-MT cell lines revealed several consistent epigenetic trends:

1. Large intergenic domains of H3K36me2 which are present in NSD1-WT cells are nearly totally absent in NSD1-MT lines.
2. DNA methylation, which is normally associated with intergenic H3K36me2 is greatly reduced in NSD1-MTs. Hence, outside of actively transcribed genes, NSD1-MT cells are globally hypomethylated.
3. The levels of the H3K27me3 modification, associated with silenced regions and antagonized by the presence of H3K36me2/3, are elevated in NSD1-MT cell lines, particularly in regions that are occupied by H3K36me2 in NSD1-WT.

Disrupting NSD1 by CRISPR-Cas9 in the three NSD1-WT cell lines allowed us to establish the extent to which these epigenomic characteristics were a direct consequence of the absence of functional NSD1. The NSD1-KO lines faithfully recapitulated the reduction of intergenic H3K36me2 and the corresponding increase in H3K27me3 observed in NSD1-MT cells. At the DNA methylation level, although DNAme decreased in the regions of H3K36me2 loss, we noted that the decrease was modest compared to the hypomethylation observed in the NSD1-MT cell lines. We also found that the extent of decrease in DNAme was variable across lines, showing that the genetic and epigenetic state of the parental cell line is an important factor in the fate of DNAme following epigenome dysregulation. The relatively small decrease in DNA methylation may be explained by the fact that, compared to histone modifications, DNA methylation is a more stable mark, and once established it tends to be more faithfully maintained, particularly in differentiated cell lines. Overall, our results strongly support the direct causal effect of NSD1 disruption on the epigenetic deregulation observed in NSD1-MT HNSCCs.

Having characterized the primary epigenetic outcomes of NSD1 deletion, we aimed to understand the downstream consequences and contributions to the pathology of HNSCC. Recent findings show that H3K36me2 helps to promote the establishment of DNA methylation^25^ and restrict the spread of heterochromatic H3K27me3^26,57^. Furthermore, H3K36me2 domains surround actively transcribed genes and are associated with “active” regions of the genome^25^. While generally not transcribed, those regions tend to be rich in chromosomal contacts, CTCF binding sites, and H3K27 acetylation peaks that are characteristic of open chromatin and CREs^39^. Our analysis demonstrates that in HNSCC the regions of NSD1-dependent H3K36me2 loss are indeed significantly enriched in CREs and specifically distal enhancers. Upon the loss of H3K36me2, those enhancers also lose DNAme, gain H3K27me3 and, most importantly, lose the active mark H3K27ac. This loss of enhancer activity is correlated with reduced expression of target genes. It will be of high mechanistic interest to understand how this loss of acetylation results from the primary epigenetic changes. It is possible that H3K36me2 is involved in promoting the activity of histone acetyltransferases. DNA methylation loss, together with loss of H3K36me2, may result in aberrant recruitment of transcription factors that are needed to enhance open chromatin state. It is also possible that chromatin compaction due to H3K27me3 spread restricts acetylation or hinders acetyltransferases by direct competition for substrates. Further studies will be needed to elucidate those questions.

Our findings on epigenetic consequences of NSD1 mutations in HNSCCs complement recent advances in understanding the significance of H3K36me2 in cancer. Lhoumaud et al.^39^ investigated the function of *NSD2* – another member of the histone methyltransferase family that has been implicated in depositing intergenic H3K36me2 – in multiple myeloma. In cells that naturally carry the 4;14 translocation that drives overexpression of *NSD2*, reducing *NSD2* levels results in depletion of intergenic H3K36me2 domains, decreased enhancer activity, and downregulation of target gene expression within topologically associated chromatin domains. In pancreatic carcinoma, Yuan et al.^51^ found opposing effects of disruption of *NSD2* and the lysine-specific demethylase *KDM2A* and concluded that the NSD2-associated reduction of H3K36me2 results in loss of enhancer activity of a specific class of enhancers that regulate EMT.

Although the downstream epigenetic effects of NSD1 and NSD2 appear similar, mutations in those two methyltransferases are involved in distinct pathologies. In cancer, overactivity of NSD2 has been implicated in blood malignancies: activating NSD2 point mutations in acute lymphocytic leukemia^58,59^, IgH-NSD2 fusion in multiple myeloma^60^. A frequent NUP98-NSD1 translocation has been found in acute myeloid leukemia^61^, although most likely this fusion does not act through NSD1 overexpression but a gain of function phenotype of the resulting protein^62^. Loss of function mutations in NSD1 has been identified as driver mutations in squamous cell carcinomas of the head and neck^1^ and lung^20^. Although some loss-of-function NSD2 mutations are found in HNSCC^63^, to our knowledge they have not been identified as statistically significant driver mutations in neither HNSCC nor any other cancer. In genetic disease, heterozygous loss of function mutations in NSD1 are responsible for Sotos overgrowth syndrome^64^, while heterozygous loss of function mutations in NSD2 have been associated with Wolf-Hirschhorn syndrome^65,66^, which is characterized by a growth deficiency. Conversely, NSD2 mutations have not been found in patients with overgrowth syndromes^67^. It is possible that the different phenotypic outcomes of mutations in NSD1 and NSD2 are a result of their different expression patterns across developmental times and tissue types, but given the differences in protein structure and numbers of alternatively spliced isoforms produced by each gene, it is likely that the two methyltransferases have other, divergent functions.

Our final aim was to understand the transcriptomic consequences of NSD1 loss in HNSCC. We carried out integrative Gene Set Enrichment Analysis, aiming to focus on the primary genes targeted by the regulatory cascade. Several pathways, including KRAS signaling, epithelial-mesenchymal transition (EMT), and inflammatory responses were downregulated following loss of NSD1 (Figure 5a). These findings further substantiate recent studies on dysregulation of H3K36me2 in other biological and disease contexts. In patients with Sotos syndrome caused by germline NSD1 haploinsufficiency, deregulation of MAPK/ERK signaling pathway downstream of KRAS activation was observed and postulated to contribute to accelerated skeletal outgrowth^52^. Similarly, NSD2-mediated H3K36me2 has been shown to contribute to KRAS transcriptional program in lung cancers^53^. NSD2 has also been implicated in promoting EMT in pancreatic carcinoma^51^, prostate cancer, renal cell carcinoma, and multiple myelomas^48-50^. A marked downregulation of immune response appeared as one of the most consistent trends across various analyses that we have conducted. This observation is in agreement with recent findings that NSD1-MT HNSCC exhibits an immune-cold phenotype with low T-cell infiltration^20,68^. It is remarkable that increasing evidence points to the association of NSD1 mutations and reduced DNAme with deficient immune response in HNSCCs, since in other cancers, such as melanoma DNA hypomethylation has been implicated with elevated immune response, possibly through de-repression of retroviral sequences and viral mimicry mechanisms^69^. Further mechanistic studies are warranted, which will have significant translational implications for the future development of immune therapies for HNSCCs.

Finally, direct cancer relevance of our principal findings, which were based on genetic manipulation of several cell lines, was demonstrated by the reanalysis of primary tumor data from TCGA HNSC. We first discovered that NSD1 truncating mutations were indeed the shared characteristic among the tumor samples whose transcriptome and methylome were most similar to the cell lines used in this study.

Further analysis revealed four trends that are consistent with our cell-line based findings:

1. DNA hypomethylation in NSD1-Mutant tumors preferentially influences CREs.
2. DNA hypomethylation in tumors is significantly associated with the regions that lost H3K36me2 as a result of the knockout of NSD1.
3. Differential gene expression comparison between NSD1-WT vs. NSD1-KO in our cell line system and NSD1-WT vs. NSD1-MT tumors showed significant similarities between downregulated genes.
4. Most of the genetic pathways – and specifically those related to immune response, EMT, and KRAS signaling – perturbed by NSD1-KO in cell culture were also similarly affected in the comparison of NSD1-WT vs. NSD1-MT tumors.

In summary, our studies characterized the extensive epigenome reprogramming induced by NSD1 loss in HNSCCs, which may, in turn, lead to multifaceted effects on tumor growth, plasticity, and immunogenicity. More work will be needed to understand why such a global chromatin perturbation, which affects much of intergenic H3K36me2, causes deregulation of specific biological pathways. Alternations in tumor lineage plasticity and immune response suggest that NSD1 could serve as a potential biomarker for patient’s response to existing chemo- or immune-therapy, respectively. Furthermore, these hallmarks may constitute vulnerabilities of the tumor that may be explored in designing therapeutic approaches.

## METHODS

### Cell culture

FaDu (ATCC), Cal27 (ATCC), Detroit 562 (ATCC), SKN-3 (JCRB Cell Bank), and SCC-4 (ATCC) cells from ATCC were cultured in Dulbecco’s modified Eagle medium (DMEM:F12; Invitrogen) with 10% fetal bovine serum (FBS; Wisent). BICR78 (ECACC) was cultured in DMEM:F12 (Invitrogen) with 10% FBS (Wisent), and 400ng/ml hydrocortisone. *Drosophila* S2 cells were cultured in Schneider’s *Drosophila* medium (Invitrogen) containing 10% heat-inactivated FBS. All cell lines tested negative for mycoplasma contamination.

### CRISPR–Cas9 gene editing and generation of stable cell lines

To generate knockout lines of Cal27, Detroit 562, and FaDu cell lines, CRISPR-Cas9 editing was performed using the Alt-R CRISPR-Cas9 System (IDT) and designing synthetic crRNA guides to form a duplex with Alt-R® CRISPR-Cas9 tracrRNA, ATTO™ 550 and coupled to the Cas9 Nuclease V3 following IDT instructions for “Cationic lipid delivery of CRISPR ribonucleoprotein complexes into mammalian cells”. Transfection was performed using Lipofectamine CRISPRMAX reagent (Thermo Fisher Scientific) with a lower volume than the company’s protocol (with the ratio of 0.05 to RNP) and Cas9 PLUS Reagent (Thermo Fisher Scientific) was used in order to improve transfection. The transfected cells were incubated for 48 h. Single ATTO550+ cells were then sorted into 96-well plates. Clones were expanded and individually verified by Sanger and MiSeq sequencing of the target loci. Guide sites sequence for NSD1 KO: guide 1 in PWWP domain: GCCCTATCGGCAGTACTACG; guide 2 in SET domain: GTGAATGGAGATACCCGTGT.

### Histone acid extraction, histone derivatization, and analysis of post-translational modifications by nano-LC–MS

Cell frozen pellets were lysed in nuclear isolation buffer (15 mM Tris pH 7.5, 60 mM KCl, 15 mM NaCl, 5 mM MgCl_2_, 1 mM CaCl_2_, 250 mM sucrose, 10 mM sodium butyrate, 0.1% v/v b-mercaptoethanol, commercial phosphatase and protease inhibitor cocktail tablets) containing 0.3% NP-40 alternative on ice for 5 min. Nuclei were washed in the same solution without NP-40 twice and the pellet was slowly resuspended while vortexing in chilled 0.4 N H_2_SO_4_, followed by 3 h rotation at 4°C. After centrifugation, supernatants were collected and proteins were precipitated in 20% TCA overnight at 4degC, washed with 0.1% HCl (v/v) acetone once and twice using acetone only, to be resuspended in deionized water. Acid-extracted histones (5–10 μg) were resuspended in 100 mM ammonium bicarbonate (pH 8), derivatized using propionic anhydride and digested with trypsin as previously described^70^. After the second round of propionylation, the resulting histone peptides were desalted using C18 Stage Tips, dried using a centrifugal evaporator and reconstituted using 0.1% formic acid in preparation for liquid chromatography-mass spectrometry (LC–MS) analysis. Nanoflow liquid chromatography was performed using a Thermo Fisher Scientific. Easy nLC 1000 equipped with a 75 µm × 20-cm column packed in-house using Reprosil-Pur C18-AQ (3 µm; Dr. Maisch). Buffer A was 0.1% formic acid and Buffer B was 0.1% formic acid in 80% acetonitrile. Peptides were resolved using a two-step linear gradient from 5% B to 33% B over 45 min, then from 33% B to 90% B over 10 min at a flow rate of 300 nl min^−1^. The HPLC was coupled online to an Orbitrap Elite mass spectrometer operating in the positive mode using a Nanospray Flex Ion Source (Thermo Fisher Scientific) at 2.3 kV. Two full mass spectrometry scans (*m*/*z* 300–1,100) were acquired in the Orbitrap Fusion mass analyzer with a resolution of 120,000 (at 200 *m*/*z*) every 8 data-independent acquisition tandem mass spectrometry (MS/MS) events, using isolation windows of 50 *m*/*z* each (for example, 300–350, 350–400…650–700). MS/MS spectra were acquired in the ion trap operating in normal mode. Fragmentation was performed using collision-induced dissociation in the ion trap mass analyzer with a normalized collision energy of 35. The automatic gain control target and maximum injection time were 5 × 10^5^ and 50 ms for the full mass spectrometry scan, and 3 × 10^4^ and 50 ms for the MS/MS scan, respectively. Raw files were analysed using EpiProfile 2.0^71^. The area for each modification state of a peptide was normalized against the total signal for that peptide to give the relative abundance of the histone modification.

### Cross linking and ChIP-sequencing

About 10 million cells per cell line were grown and directly crosslinked on the plate with 1% formaldehyde (Sigma) for 10 minutes at room temperature and the reaction was stopped using 125nM Glycine for 5 minutes. Fixed cell preparations were washed with ice-cold PBS, scraped off the plate, pelleted, washed twice again in ice-cold PBS, and flash frozen pellets stored at −80°C.

Thawed pellets were resuspended in 500ul cell lysis buffer (5 mM PIPES-pH 8.5, 85 mM KCl, 1% (v/v) IGEPAL CA-630, 50 mM NaF, 1 mM PMSF, 1 mM Phenylarsine Oxide, 5 mM Sodium Orthovanadate, EDTA-free Protease Inhibitor tablet) and incubated 30 minutes on ice. Samples were centrifugated and pellets resuspended in 500ul of nuclei lysis buffer (50 mM Tris-HCl pH 8.0, 10 mM EDTA, 1% (w/v) SDS, 50 mM NaF, 1 mM PMSF, 1 mM Phenylarsine Oxide, 5 mM Sodium Orthovanadate and EDTA-free protease inhibitor tablet) and incubated 30 minutes on ice. Sonication of lysed nuclei was performed on a BioRuptor UCD-300 at max intensity for 60 cycles, 10s on 20s off, centrifuged every 15 cycles, chilled by 4°C water cooler. Samples were checked for sonication efficiency using the criteria of 150-500bp by gel electrophoresis of a reversed cross-linked and purified aliquot. After the sonication, the chromatin was diluted to reduce SDS level to 0.1% and concentrated using Nanosep 10k OMEGA (Pall). Before ChIP reaction 2% of sonicated drosophila S2 cell chromatin was spiked-in the samples for quantification of total levels of histone mark after the sequencing.

ChIP reaction for histone modifications was performed on a Diagenode SX-8G IP-Star Compact using Diagenode automated Ideal ChIP-seq Kit. Dynabeads Protein A (Invitrogen) were washed, then incubated with specific antibodies (anti-H3K27me3 Cell Signaling Technology 9733, anti-H3K36me2 CST 2901, anti-H3K27ac Diagenode C15410196), 1.5 million cells of sonicated cell lysate, and protease inhibitors for 10 hr, followed by 20 min wash cycle using the provided wash buffers (DIAGENODE Immunoprecipitation Buffers, iDeal ChIP-seq kit for Histone).

Reverse cross-linking took place on a heat block at 65°C for 4 hr. ChIP samples were then treated with 2ul RNase Cocktail at 65°C for 30 min followed by 2ul Proteinase K at 65°C for 30 min. Samples were then purified with QIAGEN MiniElute PCR purification kit as per manufacturers’ protocol. In parallel, input samples (chromatin from about 50,000 cells) were reverse crosslinked and DNA was isolated following the same protocol. Library preparation was carried out using KAPA Hyper Prep library preparation reagents, following the manufacturer’s protocol. ChIP libraries were sequenced using Illumina HiSeq 4000 at 50bp single reads or NovaSeq 6000 at 100bp single reads.

### Whole Genome Bisulphite Sequencing

Whole genome sequencing libraries were generated from 1000 ng of genomic DNA spiked with 0.1% (w/w) unmethylated λ DNA (Roche Diagnostics) and fragmented to 300–400 bp peak sizes using the Covaris focused-ultrasonicator E210. Fragment size was controlled on a Bioanalyzer High Sensitivity DNA Chip (Agilent) and NxSeq AmpFREE Low DNA Library Kit (Lucigen) was applied. End repair of the generated dsDNA with 3’ or 5’ overhangs, adenylation of 3’ ends, adaptor ligation, and clean-up steps were carried out as per Lucigen’s recommendations. The cleaned-up ligation product was then analyzed on a Bioanalyzer High Sensitivity DNA Chip (Agilent). Samples were then bisulfite converted using the EZ-DNA Methylation Gold Kit (Zymo Research) according to the manufacturer’s protocol. DNA was amplified by 9 cycles of PCR using the Kapa Hifi Uracil + DNA polymerase (KAPA Biosystems) according to the manufacturer’s protocol. The amplified libraries were purified using Ampure XP Beads (Beckman Coulter), validated on Bioanalyzer High Sensitivity DNA Chips, and quantified by PicoGreen. Sequencing of the WGBS libraries was performed on the Illumina HiSeqX system using 150-bp paired-end sequencing.

### RNA sequencing

Total RNA was extracted from cell pellets of approximatively 1 million cells, washed with PBS, spun down and preserved at −80, using the AllPrep DNA/RNA/miRNA Universal Kit (Qiagen) according to the manufacturer’s instructions including DNase treatment option. Library preparation was performed with ribosomal RNA depletion according to the manufacturer’s instructions (NEB) to achieve greater coverage of mRNA and other long non-coding transcripts. Paired-end sequencing (100 bp) was performed on the Illumina HiSeq 4000 or NovaSeq 6000 platform.

### Western Blot

Cells are collected and counted using automatic Countess counter and 1 million cells are collected in individual test tubes and spun down. The cell pellet is washed once using PBS before spinning down again, removing the PBS, and transferring to −80degC for future use. Cell pellets are thawed on ice and resuspended in 85 to 100ul of 1x RIPA buffer from 10x (cell signaling #9806) and add 1:100 Proteinase inhibitors cocktail (P8340, Sigma) and 0.1mM of PMSF. Vortex a few times during the one-hour incubation on ice. Spin down max speed for 10 minutes at 4degC. Collect the supernatant to new tubes and proceed to quantification using BCA-Pierce Protein assay ThermoScientific/Pierce (23227) and use 96 cell microplate flat bottom with 25ul standards every time and 5ul protein samples. Read on Infinite 200Pro Tecan-icontrol. All from Bio-Rad, use stain-free TGX 4-15% gradient pre-cast gels (4568084), 1x Tris-Glycine running buffer (1610732), for protein standards mix equal amount of all-blue (1610373) and unstained (1610363). As laemmli, use premade 6x buffer containing 0.35M Tris HCl pH 6.8, 30% Glycerol, 10% SDS, 20% Beta-mercaptoethanol, 0.04% Bromophenol blue, in water. Load 50 ul of samples, or standards, or Laemmli 1x in each well. Run for 2/3-3/4 of the gel. Transfer using Bio-Rad trans-blot Turbo Transfer system and the included PVDF membrane in RTA kit low fluorescence (1704274) according to manufacturer instructions and transfer at the High MW program for 10min. The gel was cross-linked on Bio-rad imager system and whole protein images were captured on both gel and membranes. Blocking 1h in 5% skim milk (SM) and overnight incubation rotating at 4degC using 1ug/ml NeuroMab anti-NSD1 N312/10 sold by Antibodies Inc. (75-280) in 2% SM diluted in TBS-tween 0.1% (TBSt). Three washes of 5 minutes each on rotator were done using TBSt before and after the 1h incubation of the membranes with 1:1000 goat anti-mouse-HRP (Jackson Immunoresearch, 115-035-003) in 2%SM in TBSt. ECL Clarity (1705060) or Clarity Max (1705062) from BioRad are used to image the protein.

### Visualization

Unless otherwise stated, figures were created using ggplot2^72^ v3.3.0 or matplotlib^73^ v3.2.1. Coverage/alignment tracks were visualized using pyGenomeTracks^74^ v3.2.1 or IGV^75^ v2.8.2. Sequence logos were generated using ggseqlogo^76^ v0.1.

### Processing of sequence data

Sequences were all aligned to the GRCh38 analysis set. Reads from ChIP-seq and targeted sequencing for knock-out validation were mapped using BWA^77^ v0.7.17 with default settings of the BWA-MEM algorithm. WGBS reads were adapter and quality (Q10) trimmed using BBDuk from BBTools v38.73 (https://sourceforge.net/projects/bbmap/) (t=10 ktrim=r k=23 mink=11 hdist=1 tpe tbo qtrim=rl trimq=10 minlen=2) and aligned as well as deduplicated using BISCUIT v0.3.12 (https://github.com/zhou-lab/biscuit) with default options. Per-base methylation calling was performed with MethylDackel v0.4.0 (https://github.com/dpryan79/MethylDackel) after excluding biased ends. RNA-seq reads were aligned using STAR^78^ v2.7.3a based on GENCODE^79^ Release 33 annotations with the ENCODE standard options. Gene expression quantification was performed via Salmon^80^ v1.1.0 using default settings of the gentrome-based option. ENCODE blacklisted regions^81^ were excluded from all analyses. Variants were identified with GATK v4.1.5.0 using HaplotypeCaller.^82^

### ChIP-seq analysis

Raw tag counts were binned into windows using bedtools^83^ v2.29.0 with intersectBed (-c) in combination with the makewindows command. Library size normalization consisted of dividing binned tag counts by the total number of mapped reads after filtering, while input normalization involved taking the log2 ratio of ChIP signals by those of the input (i.e., without immunoprecipitation) with the addition of pseudocount (1) to avoid division by 0. Additionally, quantitative normalization entailed the multiplication of original signal (either in CPM or as log2 ratio over input) by the genome-wide modification percentage information obtained from mass spectrometry.

Enrichment matrices for aggregate plots and heatmaps were generated through deepTools^74,84^ v3.3.1 using bamCoverage/bamCompare (--skipZeroOverZero --centerReads --extendReads 200) followed by computeMatrix (scale-regions --regionBodyLength 20000 --beforeRegionStartLength 20000 -- afterRegionStartLength 20000 --binSize 1000). Genic regions were taken as the union of any intervals having the “gene” annotations in Ensembl, and intergenic regions were thus defined as the complement of genic ones. The ratio of intergenic enrichment over neighboring genes was calculated by dividing the median CPM of intergenic bins over the median of flanking genic bins after excluding the 10 bins near boundaries (i.e., TSS/TES) to eliminate edge effects and the outer 5 genic bins on each end to keep a comparable number of bins between genic and intergenic regions.

Unless otherwise stated, input-normalized enrichment in windows was used for analyses based on 10kb binned signals. Bins depleted in signal across all tracks (i.e., raw read count consistently lower than 100 in 10 kb bins) were excluded from further analyses. Identification of similarly behaving bin clusters were performed using HDBSCAN^85^ v0.8.24 with identical parameters for all samples (minPts = 5000, eps = 5000), and the intersection of label assignments were taken for pairwise comparisons between individual samples of the two conditions to be compared.

Overlap enrichment was determined with all the bins as the background set as implemented in LOLA^86^ v1.16.0 for Ensembl^32^ 97 annotations (genes and regulatory build^33^). Intergenic or genic ratio for quantiles (as in the microplots along the diagonal in Fig. 2e) or groups of bins (as in the hexagonal clumping in the middle panel of Fig. 3a) was computed by taking the ratio between the number of 10 kb bins completely overlapping annotated genes and those that fall entirely outside.

Enhancer annotations (double-elite) were obtained from GeneHancer^38^ v4.14. H3K27ac peaks were called using MACS^87^ v2.2.6 (-g hs -q 0.01). Differentially bound peaks were identified using DiffBind v2.14.0 (https://doi.org/10.18129/B9.bioc.DiffBind). Distribution across gene-centric annotations was obtained using ChIPseeker^88^ 1.22.1, whereas peak distance relative to TSSs was determined based on refTSS^89^ v3.1. Differential motif activity was determined using GimmeMotifs^90^ v0.14.3 with maelstrom and input being differentially bound sites labeled as either up- or down-regulated against a database of clustered motifs with reduced redundancy (gimme.vertebrate.v5.0). Motif density was calculated using HOMER^91^ v4.11 with annotatePeaks (-hist 5).

### WGBS analysis

Methylation calls were binned into 10kb windows, with per-window beta values calculated as (# methylated reads in bin) / (total # of reads in bin). Unless otherwise stated, such tracks were treated identically as ChIP-seq for analyses involving both assays. Differential methylation within actively transcribed regions was based on the union of active genes. Compatibility between cell line samples and TCGA tumors was determined by constructing a matrix of beta values for CpG sites included in Illumina 450k and is well-covered by WGBS. Correlation between columns (i.e., samples) of this matrix was then computed, enabling the subsequent computation of average Spearman correlation for a given tumor sample to all cell line samples within each condition. The relative similarity metric is finally defined as the average correlation to KO samples subtracted from those to WT.

### Hi-C analysis

TADs were identified on the merged A549 replicates using SpectralTAD^92^ v.1.2.0 allowing for 3 levels.

### RNA-seq analysis

Differential gene expression analyses were performed using DEseq2^93^ v1.26.0. Adjusted log fold changes (LFC) were calculated using apeglm^94^ v1.8.0. Significantly differentially expressed genes were selected with a s-value (null hypothesis being |adjusted LFC| < 0.5) threshold of 0.05. Significance of consistency between NSD1-WT vs KO and NSD1-WT vs MT was evaluated using RRHO2^54^ v1.0 with hypergeometric testing and stratified (split) presentation. Active genes were identified using zFPKM^27^ v1.8.0 with a threshold of −3. Rank aggregation was performed using RobustRankAggreg^95^ v1.1 with aggregateRanks (method=RRA). Gene set enrichment analyses were performed using fgsea^96^ v1.12.0 with fgseaMultilevel (minSize = 15, maxSize = 500, absEps = 0.0) against MSigDB^47^ v7.1. Relative similarity between TCGA samples and cell lines are computed in a similar manner as previously described for DNA methylation, with the exception that a gene expression matrix substituted the beta value matrix.

## STATISTICAL CONSIDERATIONS

Enrichment testing was performed using one-sided Fisher’s exact test of enrichment unless otherwise stated. P-values were converted to symbols through: 0 “***” 0.001 “**” 0.01 “*” 0.05 “?” 0.1 “” 1. Logistic regression was performed using a generalized linear model as implemented in the R stats v3.6.1 package. Differences between NSD1-WT and MT samples involved first averaging within conditions whereas those between NSD1-WT and KO involved subtracting within lines before averaging across. Unless otherwise stated: Cal27-KO corresponds to replicate 1, Det562-KO to replicate 2, and FaDu-KO to replicate 1.

## DATA AND CODE AVAILABILITY

### CODE AVAILABILITY

Custom scripts are available upon request.

### DATA AVAILABILITY

All other relevant data supporting the key findings of this study are available within the article and its Supplementary Information files or from the corresponding author upon request. Scripts used to generate results and figures are available through GitHub. Source data are provided as Supplementary Data files. Any additional source data will be provided upon request.

## ACKNOWLEDGMENTS

The work in J.M.’s lab is supported by the Large-Scale Applied Research Project grant Bioinformatics and Computational Biology grant from Genome Quebec, Genome Canada, the Government of Canada and the Ministère de l’Économie, de la Science et de l’Innovation du Québec; and NIH grant P01-CA196539. The work in B.A.G.’s lab is supported by NIH Grants R01AI118891; P01CA196539; and Leukemia and Lymphoma Robert Arceci Scholar award. C.L. is supported by NIH grant R00CA212257 and Pew-Stewart Scholars for Cancer Research award. B.H. is supported by studentship awards from the Canadian Institutes of Health Research and the Fonds de Recherche Québec – Santé. Computational analysis was performed using infrastructure provided by Compute Canada and Calcul Quebec.

## AUTHOR CONTRIBUTIONS

N.F., C.H., B.H and J.M. conceived and designed the study. N.F. and C.H. carried out the laboratory experiments. B.H. designed and carried out the computational analysis of the data with some guidance and supervision from JM. E.B. performed data pre-processing and adapted bioinformatics pipelines for analyses. X.C., Y.L. and C.L. shared their early results and provided the guidance necessary to initiate and shape this study. M.C. performed quantitative mass spectrometry analyses under the supervision of B.A.G. N.F., C.H., B.H and J.M jointly wrote the manuscript.

## ADDITIONAL INFORMATION

Supplementary Information accompanies this paper at https://www.cell.com/cell-genomics

### Competing Interests

The authors declare no competing interests.

---

**Supplementary Figure 1.**
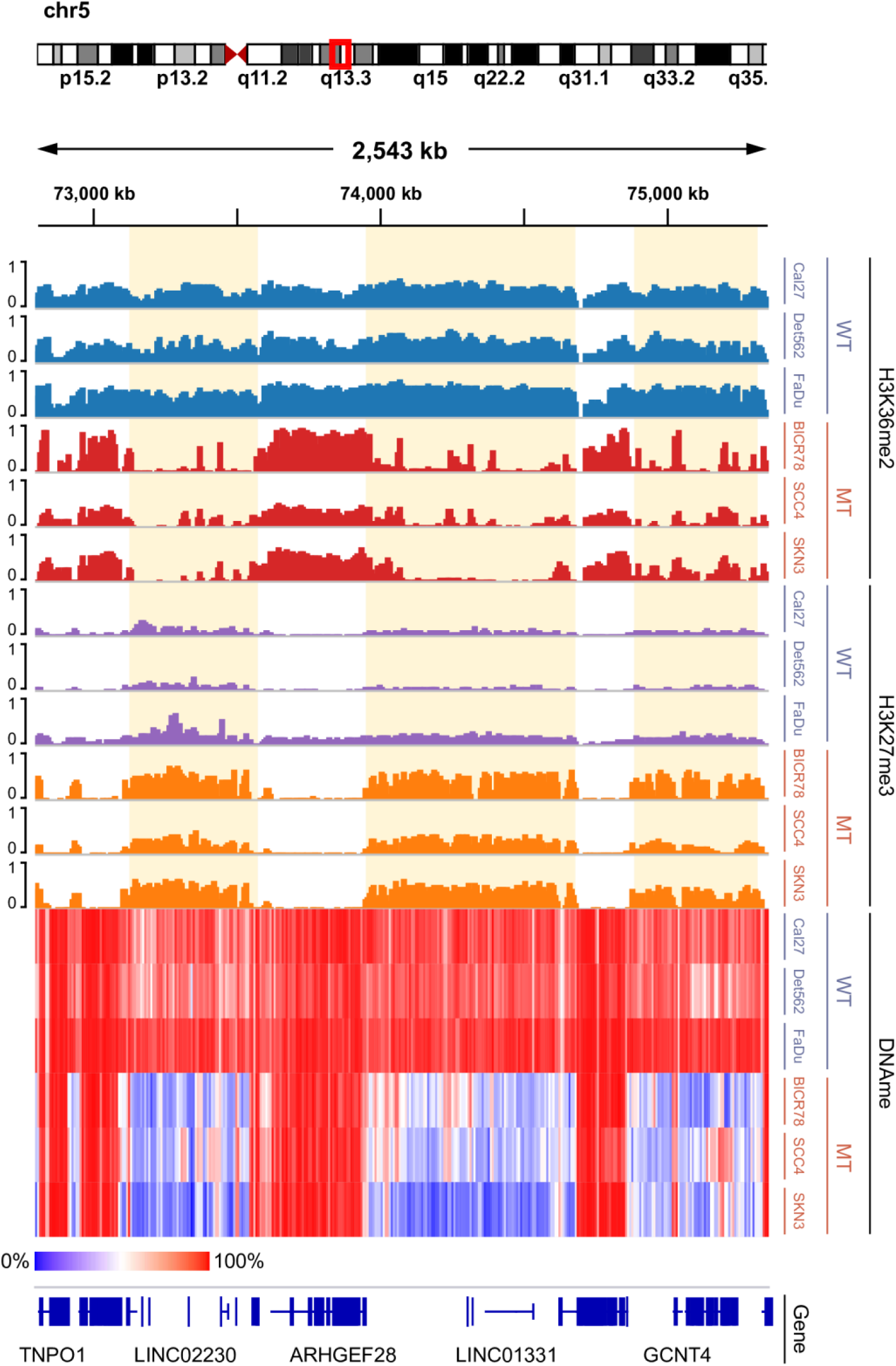
IGV screen of individual tracks making up figure 1b. Genome-browser tracks displaying individual samples; ChIP-seq signals shown are MS-normalized logCPM while beta values are used for WGBS heatmap.

**Supplementary Figure 2.**
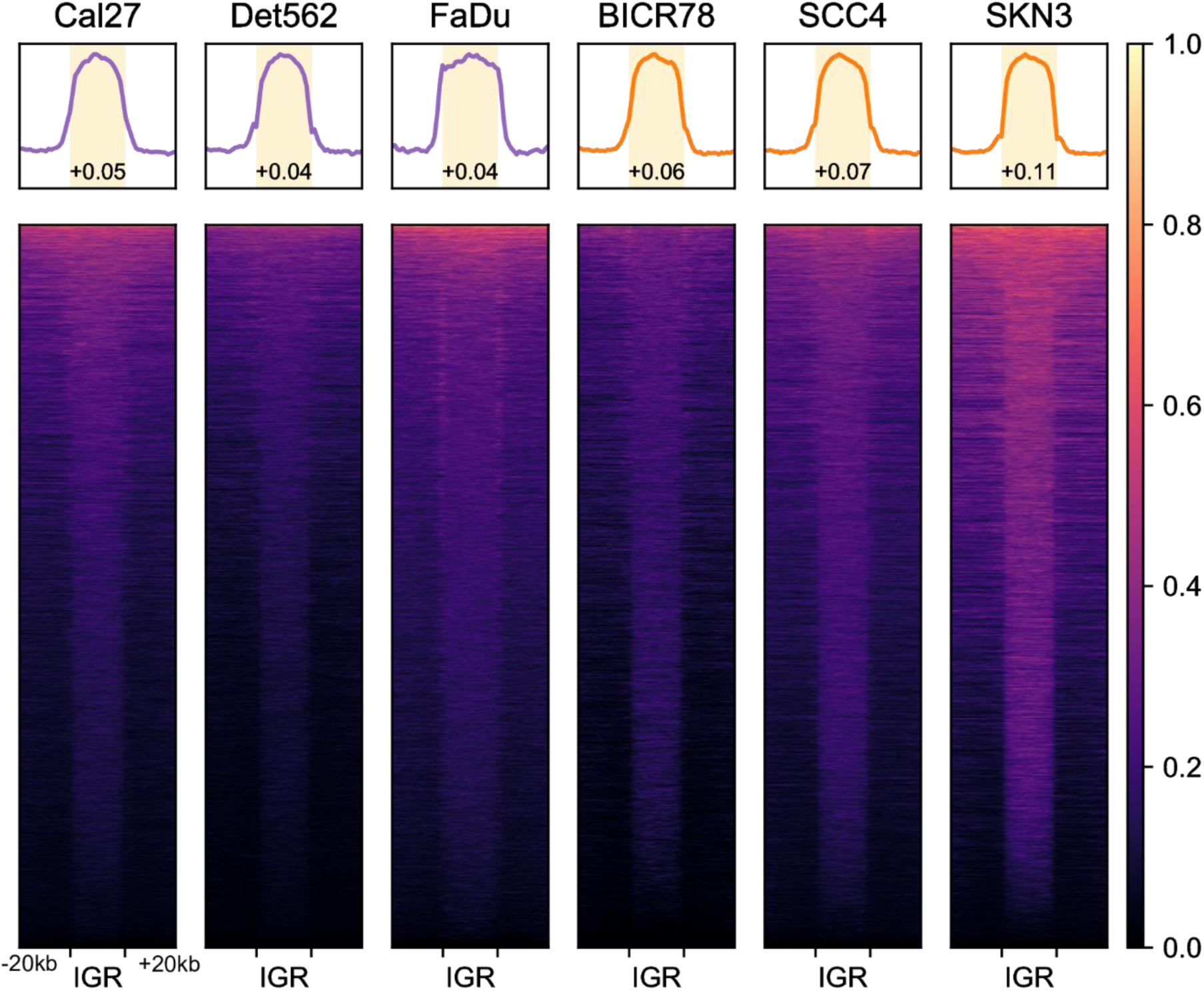
Intergenic H3K27me3 heatmap of NSD1-WT and MT samples. Heatmaps showing H3K27me3 (MS-normalized logCPM) enrichment patterns near intergenic regions. Number displayed at the bottom of aggregate plots correspond to the intergenic / genic ratio where TSS/TES and outer edges are excluded. Further details can be found the methods section.

**Supplementary Figure 3.**
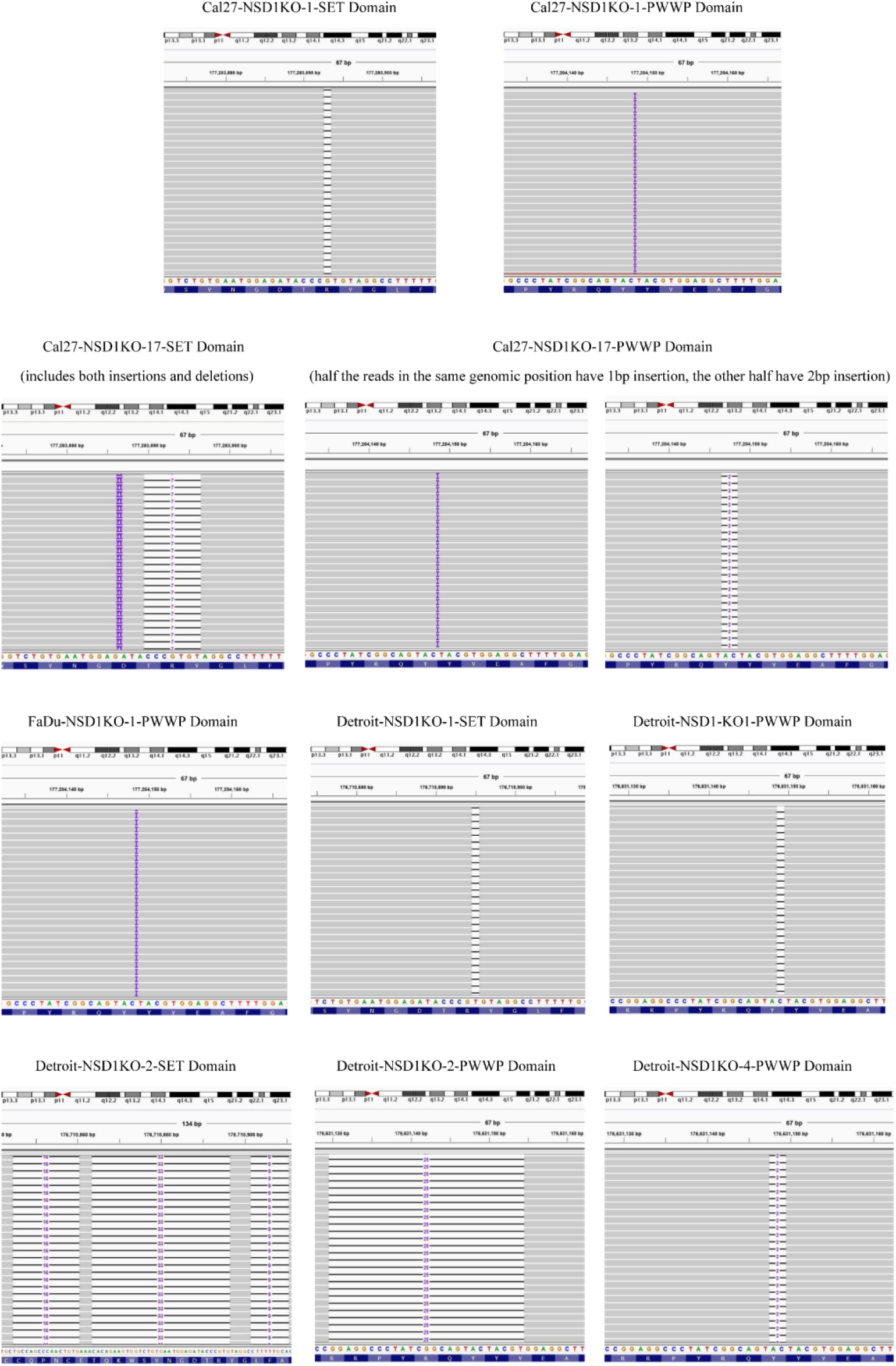
Confirmation of CRISPR-Cas9 editing. IGV Representation of Miseq data confirming the homozygous, out-of-frame mutations resulted from CRISPR-Cas9 editing in different NSD1-KO clones of Cal27, FaDu, and Detroit562 cell lines.

**Supplementary Figure 4.**
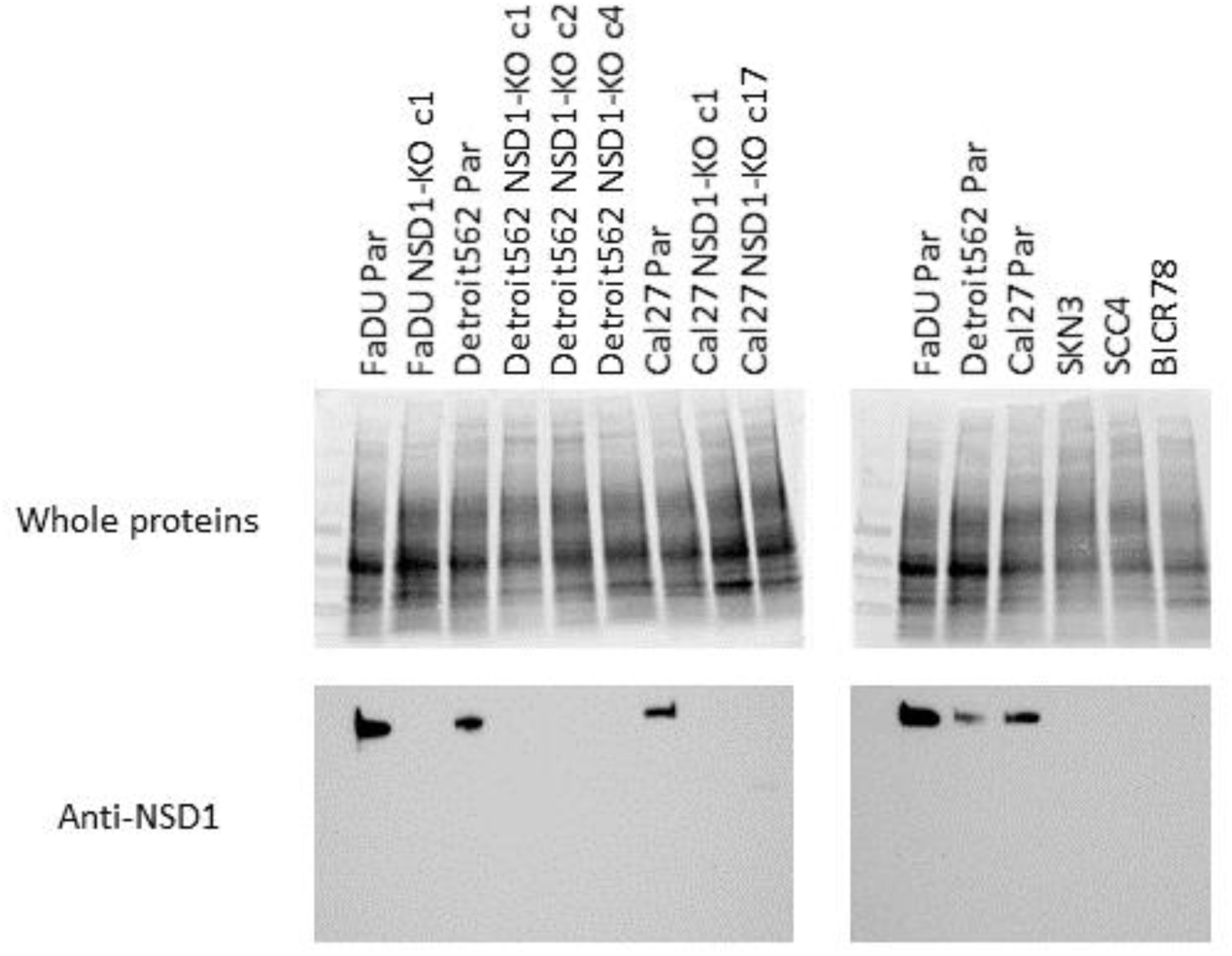
Western blots of NSD1 using 45ug protein lysates per lane, showing removal of NSD1 in all the NSD1-KO clones generated from different cell lines (Cal27, FaDu, Detroit562). Stain-free membranes showing cross-linked whole proteins after transfer are represented (top) as well as NSD1 blot (below). Expected band is at 300 kDa. The marker is Precision Plus Protein All Blue Prestained Standards.

**Supplementary Figure 5.**
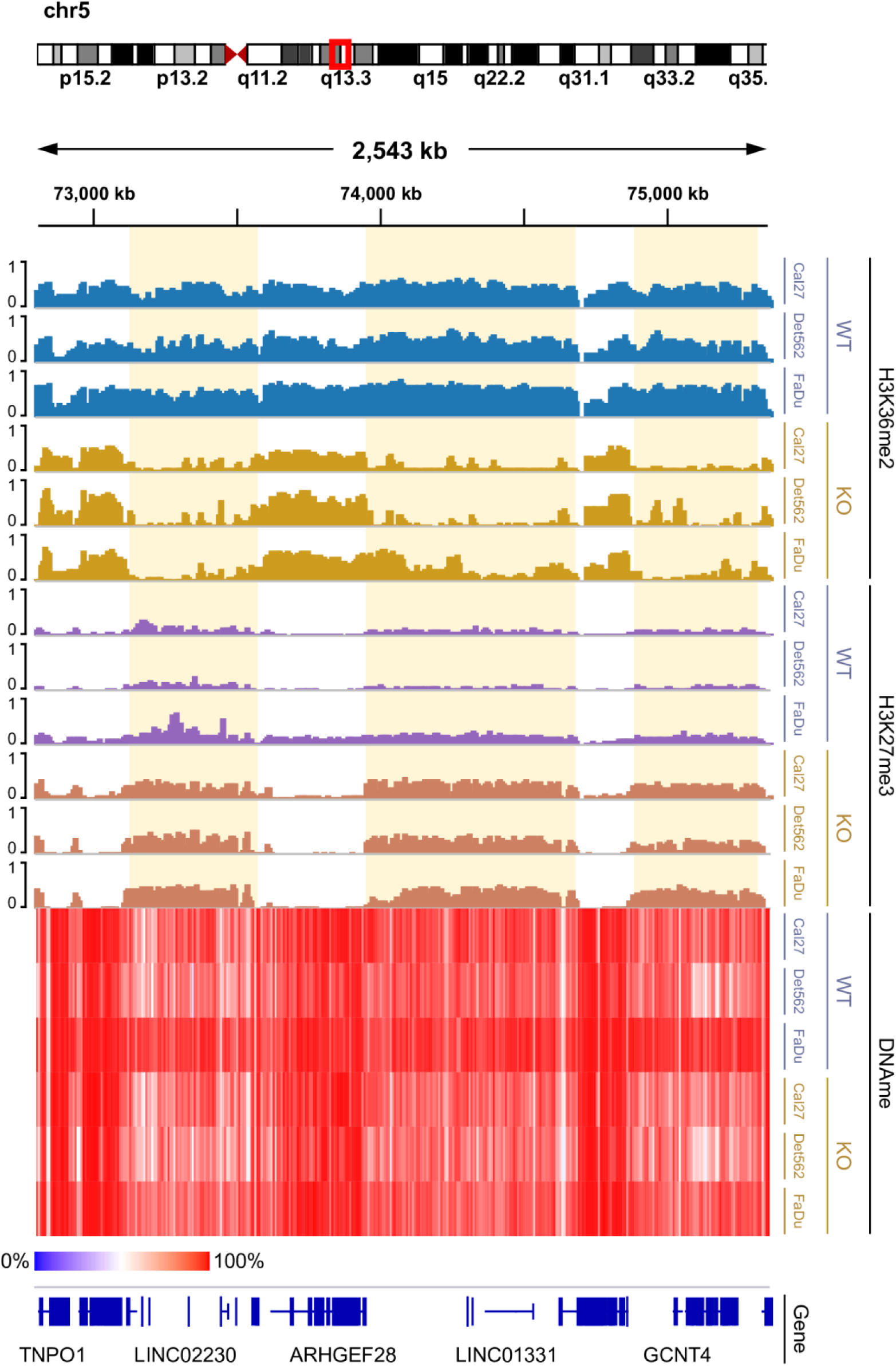
IGV screen of individual tracks making up figure 2b. Genome-browser tracks displaying individual samples; ChIP-seq signals shown are MS-normalized logCPM while beta values are used for WGBS heatmap.

**Supplementary Figure 6.**
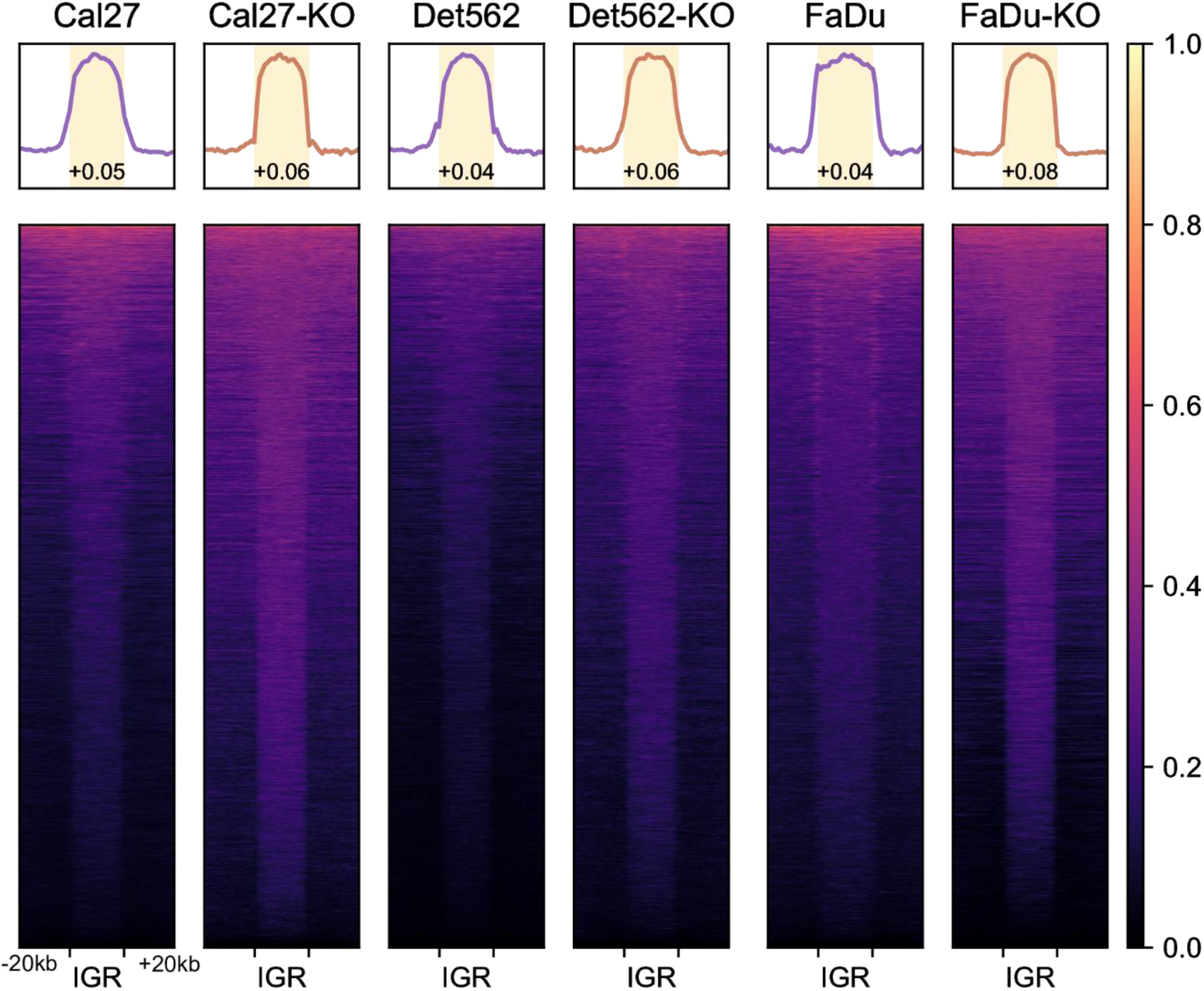
Intergenic H3K27me3 heatmap of NSD1-WT and KO samples. Heatmaps showing H3K27me3 (MS-normalized logCPM) enrichment patterns near intergenic regions. Number displayed at the bottom of aggregate plots correspond to the intergenic / genic ratio where TSS/TES and outer edges are excluded. Further details can be found the methods section.

**Supplementary Figure 7.**
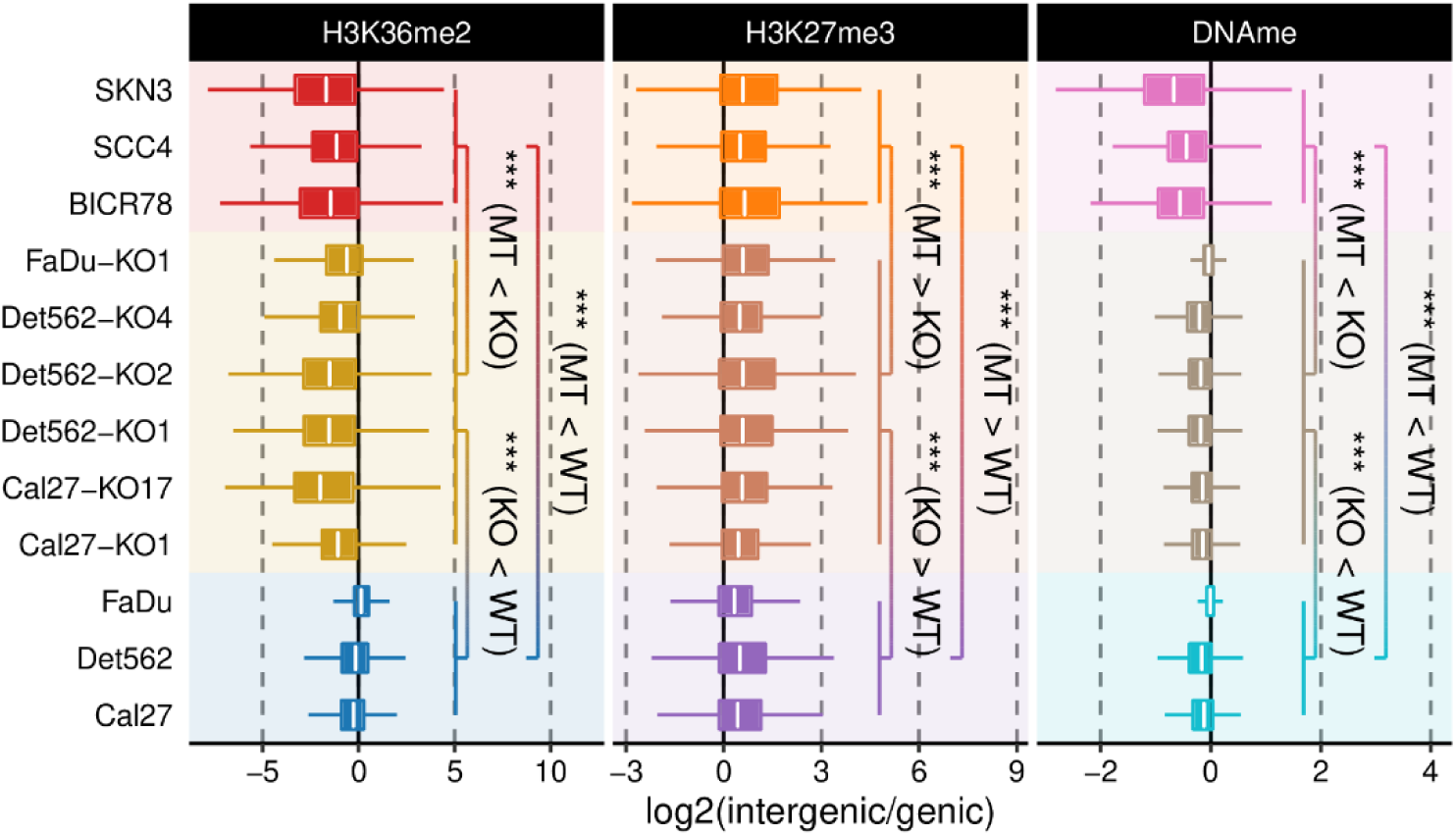
Relative enrichment of signal within intergenic regions over those of flanking genes. Intergenic enrichment/depletion relative to flanking genes to quantify the depth of “dip” or “bump” observed in heatmaps. TSS/TES and outer edges are excluded when deriving intergenic / genic ratios. Further details can be found in the methods section.

**Supplementary Figure 8.**
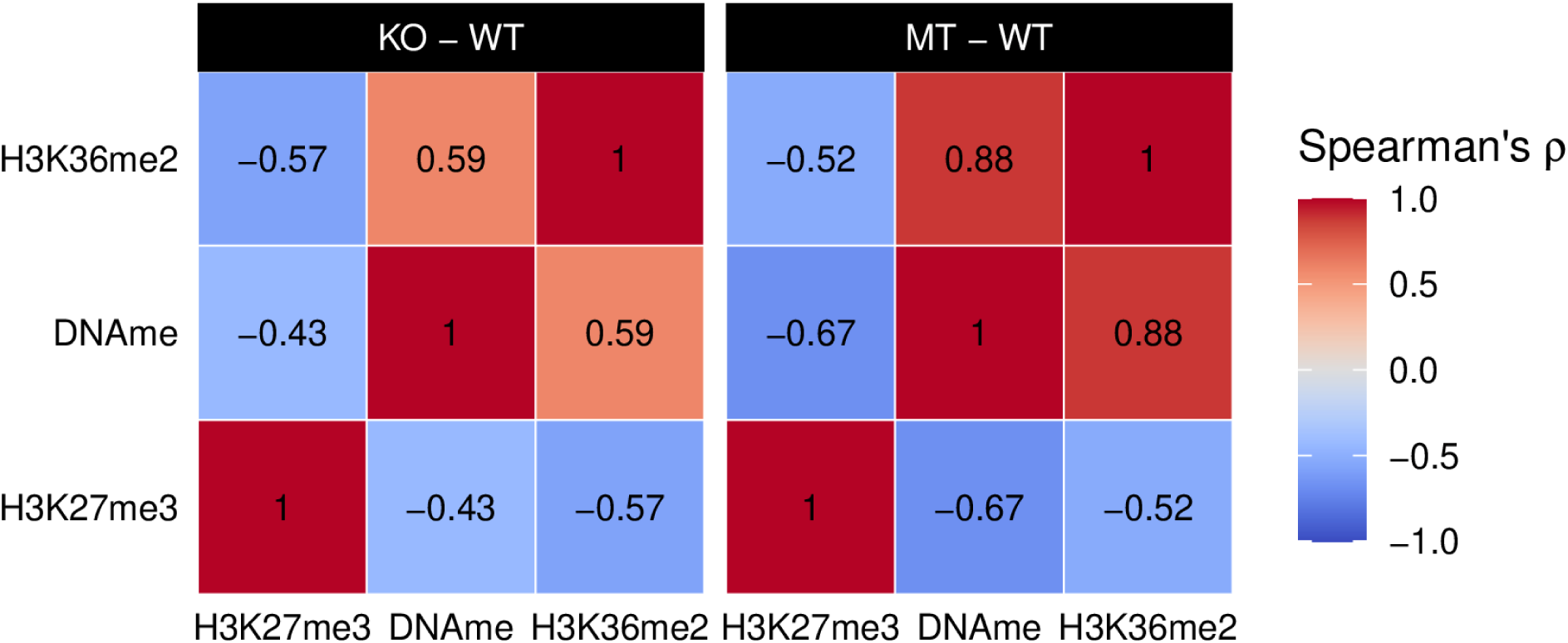
Correlation across marks in KO-WT and MT-MT comparisons. Spearman correlation matrix of differential enrichment between various combinations of marks. MT – WT comparisons involve averaging within condition before taking the difference, while for KO – WT each KO’s parental signal was subtracted before averaging across cell lines.

**Supplementary Figure 9.**
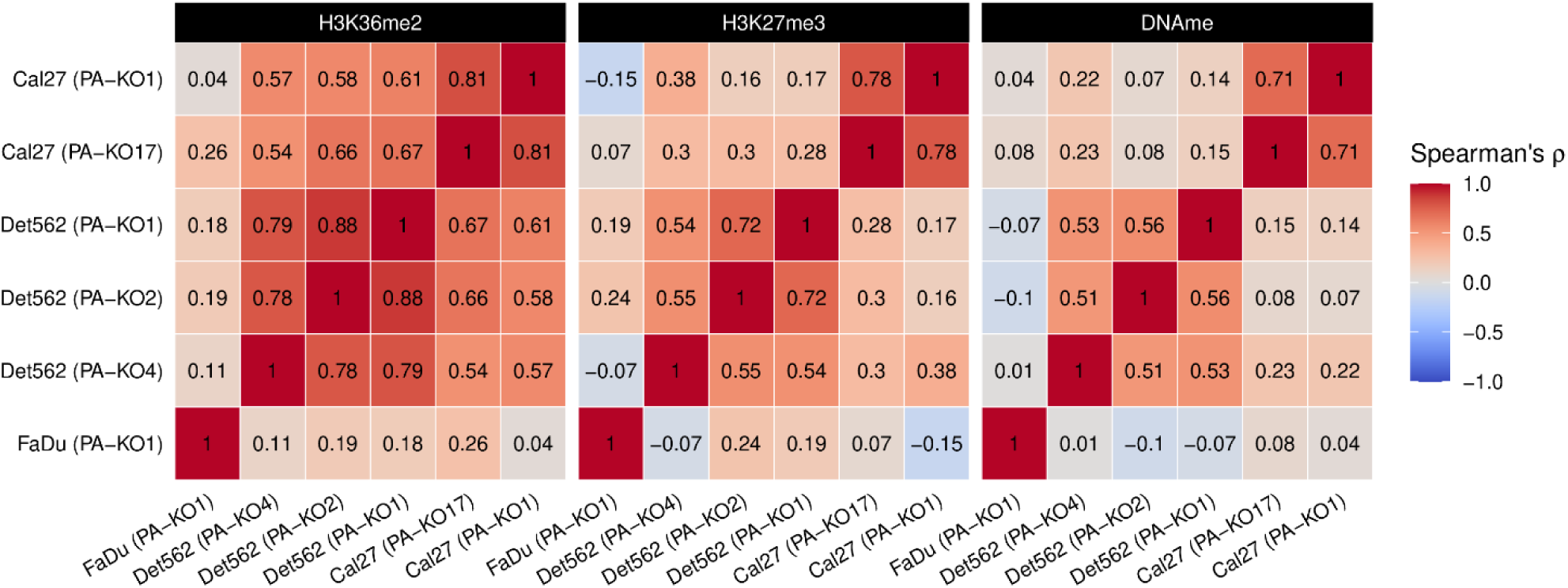
Consistency of NSD1-KO’s epigenetic impact. Spearman correlation matrix of differential enrichment for each mark between various KO-PA pairs.

**Supplementary Figure 10.**
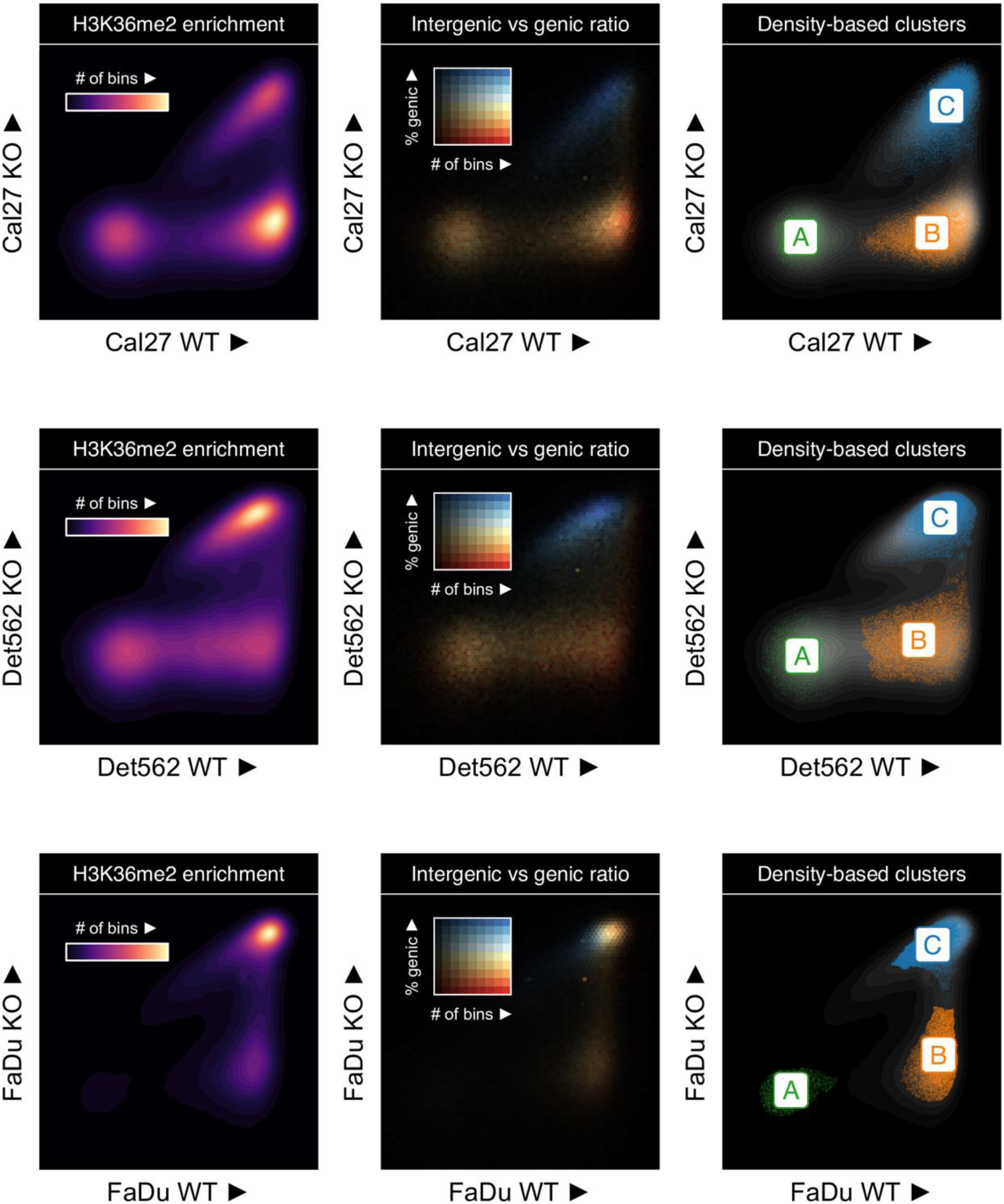
Clustering of 10 kb bins based on H3K36me2 differences. Scatterplots of H3K36me2 enrichment in 10kb windows comparing a representative WT parental samples against their NSD1-KO counterparts

**Supplementary Figure 11.**
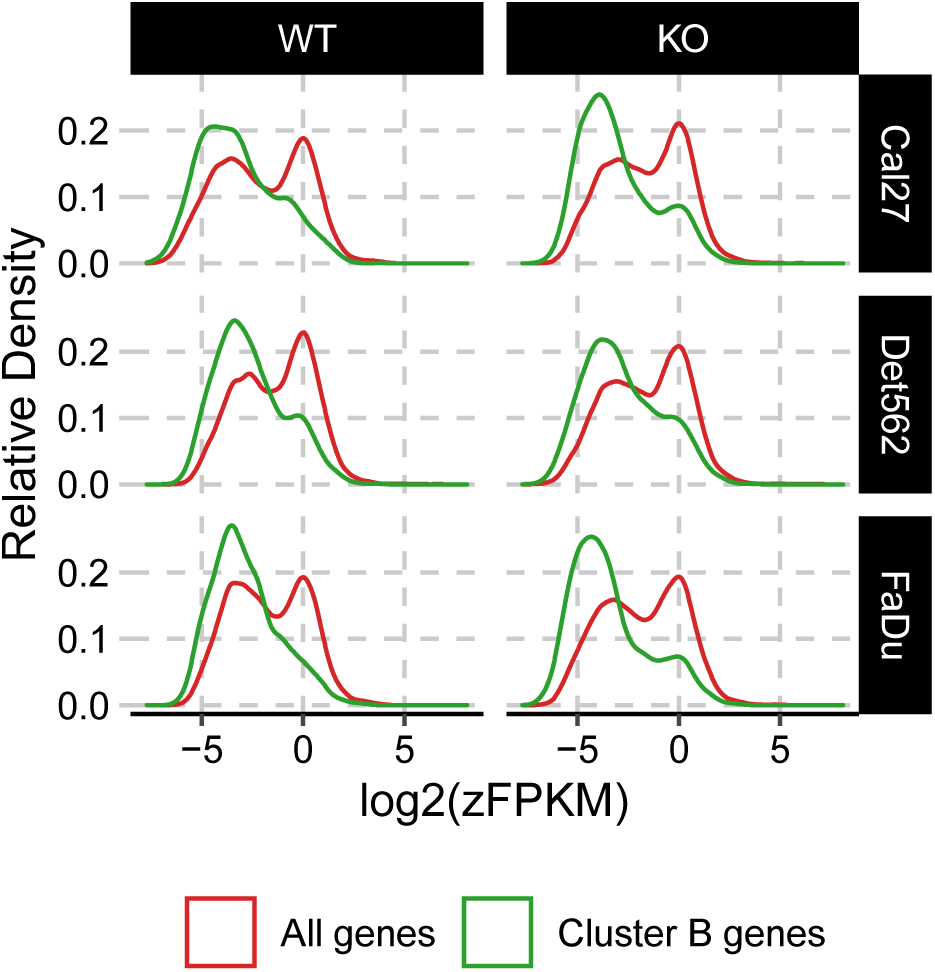
Genes overlapping cluster B bins are lowly expressed. zFPKM normalized expression level of all genes or those overlapping cluster B bins are shown

**Supplementary Figure 12.**
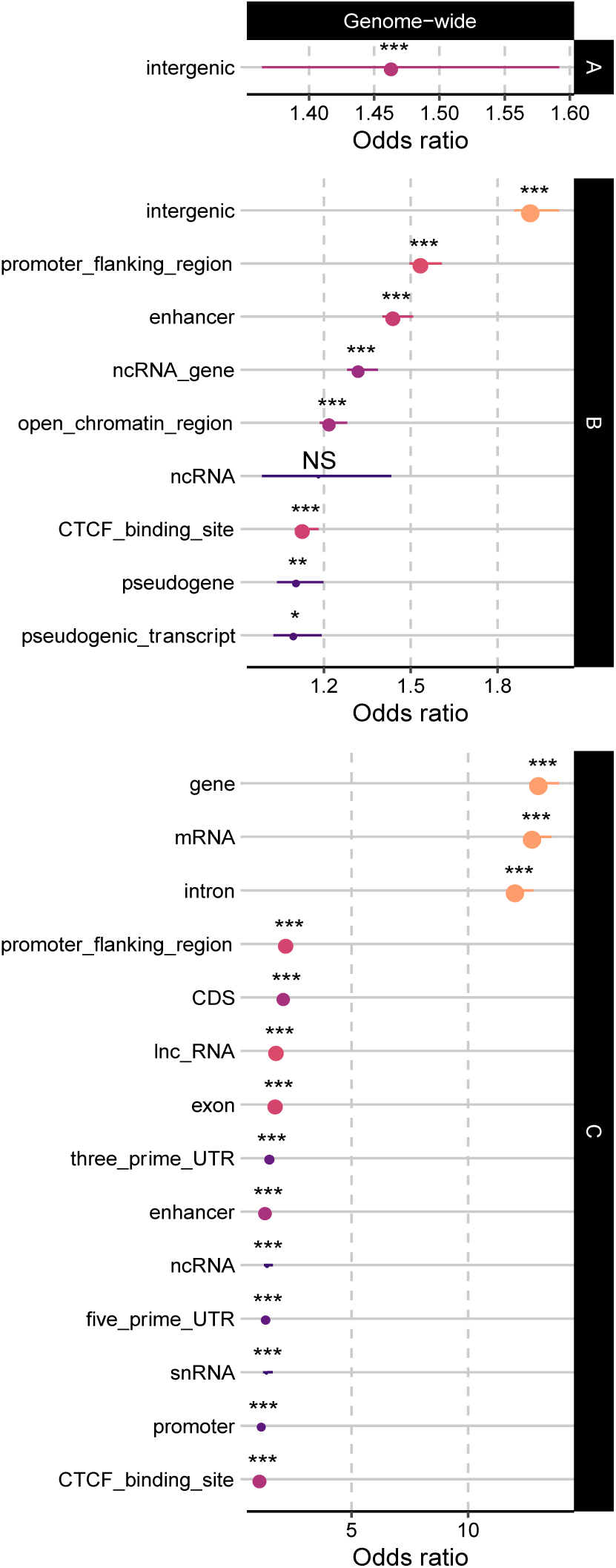
Associated annotations of bin clusters described in figure 3a. Results from overlap enrichment analysis of bins in consensus cluster A/B/C (i.e., consistently identified as a particular label for all WT vs KO comparisons) against Ensembl annotations, against a genome-wide background.

**Supplementary Figure 13.**
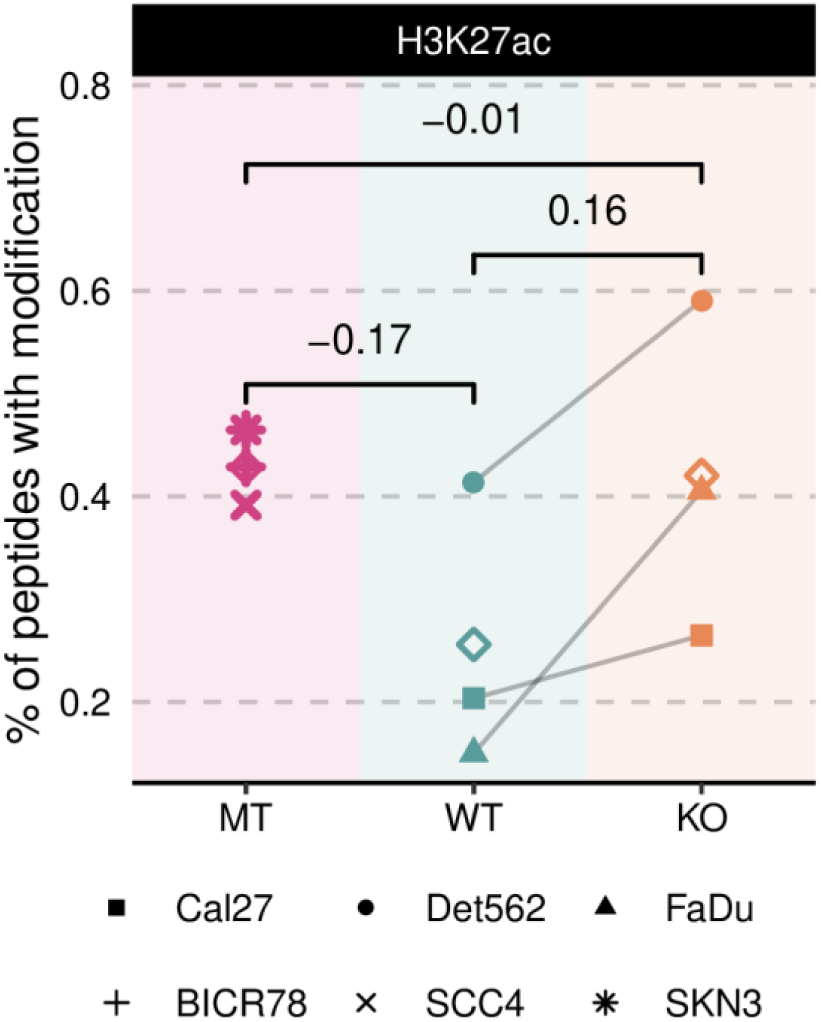
Mass spectrometry results for H3K27ac. Genome-wide prevalence of modifications based on mass spectrometry

**Supplementary Figure 14.**
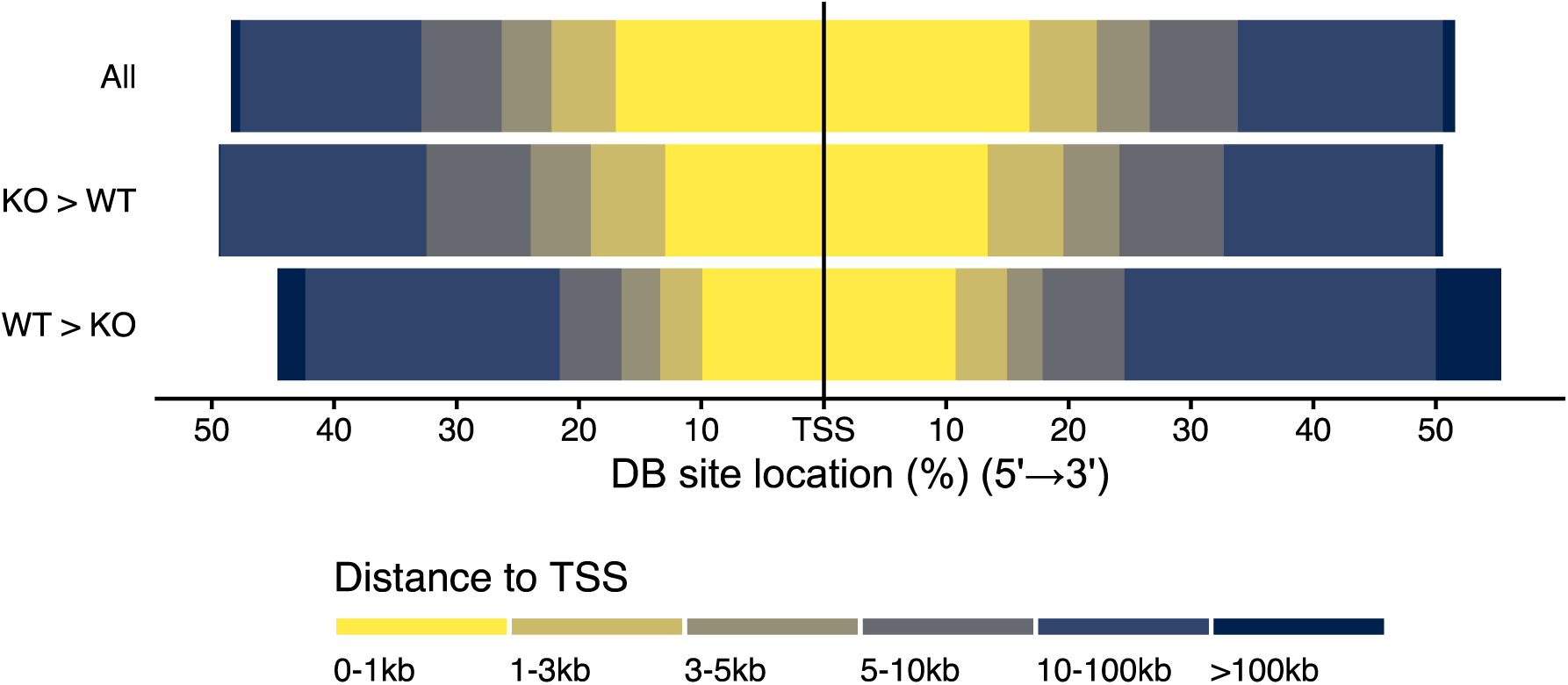
Down-regulated H3K27ac sites are more intergenic. Position of H3K27ac peaks in different classes relative to transcription start sites

**Supplementary Figure 15.**
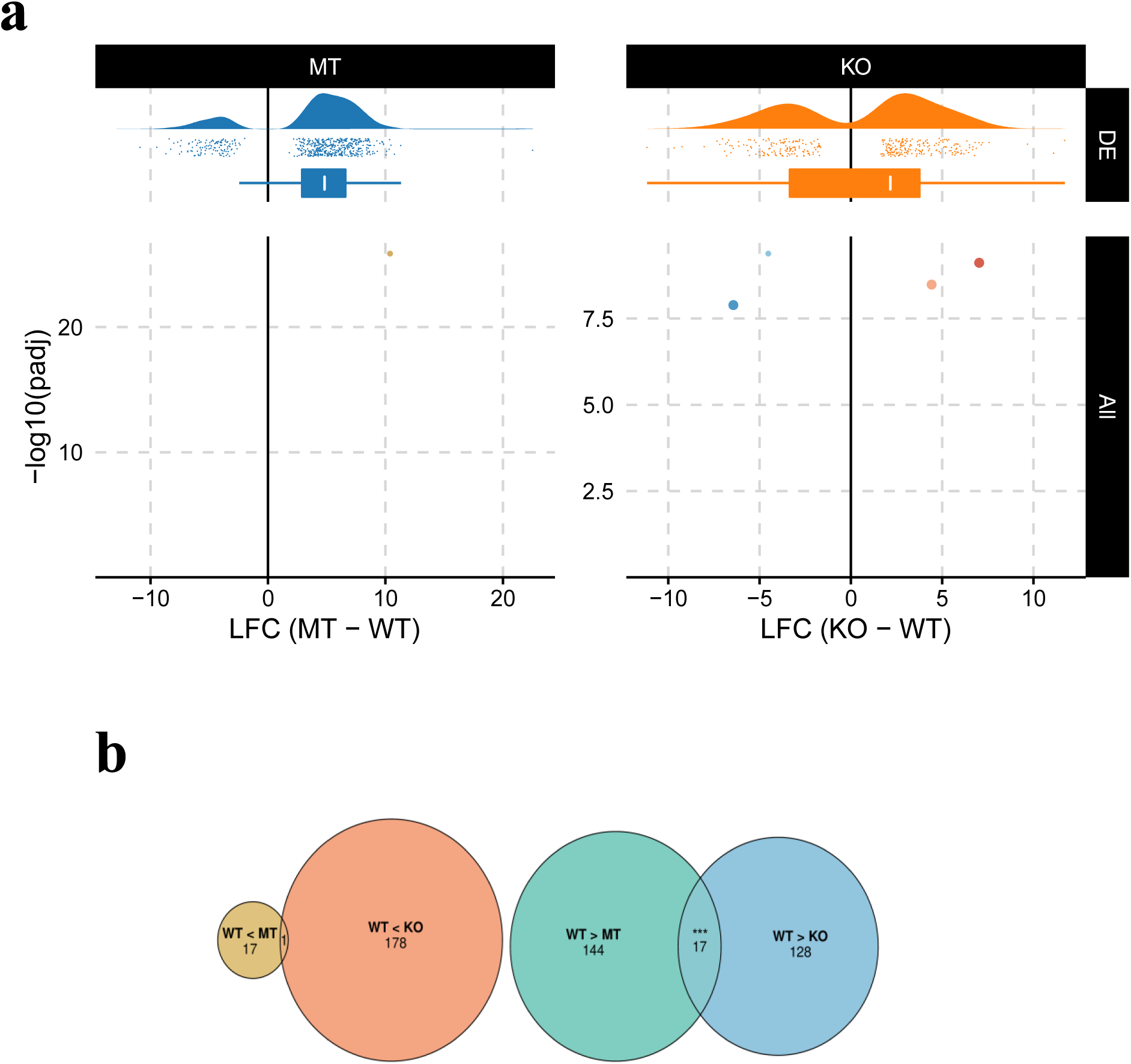
Differential gene expression analysis. **a** Volcano plot of differential gene expression magnitude and significance (bottom) and distribution of fold change for significant ones (top). **b** Overlap of differentially expressed genes between NSD1-WT vs MT and vs K0 cell line comparisons

**Supplementary Figure 16.**
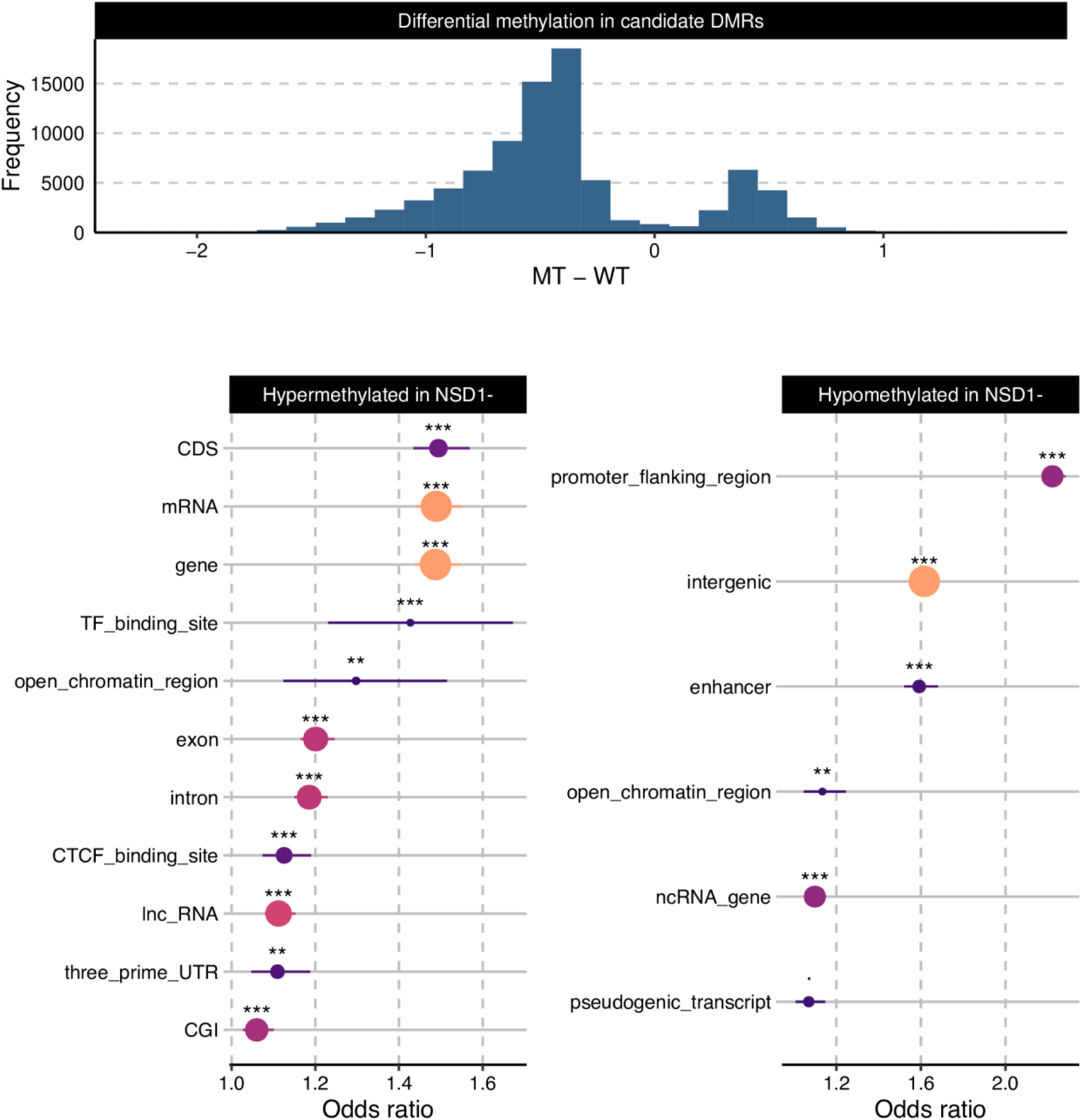
Regional specificity of NSD1-driven DNA hypomethylation. **a** CpG sites with p-value < .05 from EWAS were formed into candidate DMRs, whose NSD1 mutation status effect size are shown. **b** Results from overlap enrichment analysis of CpG sites in significantly hyper/hypomethylated DMRs against Ensembl annotations, using the set of all probes as the background.

## Supplementary Tables

**Supplementary Table 1.** Information regarding the cell lines used in this study.

| Sample Name | Disease | Age | Gender | Tissue of origin | NSD1 state |
| --- | --- | --- | --- | --- | --- |
| Cal27 | squamous cell<br>carcinoma | 56 | male | tongue | wildtype |
| FaDu | squamous cell<br>carcinoma | 56 | male | pharynx | wildtype |
| Detroit 562 | squamous cell<br>carcinoma | adult,<br>exact age<br>unspecified | female | pharynx | wildtype |
| SCC-4 | squamous cell<br>carcinoma | 55 | male | tongue | mutant |
| SKN-3 | squamous cell<br>carcinoma | unspecified | unspecified | oral cavity | mutant |
| BICR 78 | squamous cell<br>carcinoma | unspecified | unspecified | oral alveolus | mutant |

Supplementary Table 2.

Variant calls from targeted MiSeq of NSD1 locus to validate successful edits in HPV(-) NSD1-WT HNSCC cell lines used in this study.

Supplementary Table 3.

GSEA enrichment results of pathways associated with the aggregated ranking of test statistics from differential gene expression and differential acetylation of associated cis-regulatory elements in parental vs NSD1-KO cell line comparisons.

Supplementary Table 4.

GSEA enrichment results of pathways associated with the aggregated ranking of adjusted differential gene expression in both NSD1-WT vs MT and vs KO cell line comparisons.

## Supplementary Data

Supplementary Data 1.

Genome-wide prevalence of histone H3 modifications based on quantitative mass spectrometry

Supplementary Data 2.

Results from overlap enrichment analysis of bins in consensus cluster A/B/C (i.e., consistently identified as a particular label for all WT vs KO comparisons) against Ensembl annotations

Supplementary Data 3.

Summary of logistic regression models for the spatial relationship between H3K36me2 depletion and decreased expression

## Notes

### Competing Interest Statement

The authors have declared no competing interest.

### Summary of Updates

A section validating cell line results in TCGA tumor data was added to the manuscript.

